# FecalSeq: methylation-based enrichment for noninvasive population genomics from feces

**DOI:** 10.1101/032870

**Authors:** Kenneth L. Chiou, Christina M. Bergey

## Abstract

Obtaining high-quality samples from wild animals is a major obstacle for genomic studies of many taxa, particular at the population level, as collection methods for such samples are typically invasive. DNA from feces is easy to obtain noninvasively, but is dominated by a preponderance of bacterial and other non-host DNA. Because next-generation sequencing technology sequences DNA largely indiscriminately, the high proportion of exogenous DNA drastically reduces the efficiency of high-throughput sequencing for host animal genomics. In order to address this issue, we developed an inexpensive methylation-based capture method for enriching host DNA from noninvasively obtained fecal DNA samples. Our method exploits natural differences in CpG-methylation density between vertebrate and bacterial genomes to preferentially bind and isolate host DNA from majority-bacterial fecal DNA samples. We demonstrate that the enrichment is robust, efficient, and compatible with downstream library preparation methods useful for population studies (e.g., RADseq). Compared to other enrichment strategies, our method is quick and inexpensive, adding only a negligible cost to sample preparation for research that is often severely constrained by budgetary limitations. In combination with downstream methods such as RADseq, our approach allows for cost-effective and customizable genomic-scale genotyping that was previously feasible in practice only with invasive samples. Because feces are widely available and convenient to collect, our method empowers researchers to explore genomic-scale population-level questions in organisms for which invasive sampling is challenging or undesirable.

## INTRODUCTION

The past decade has witnessed a rapid transformation of biological studies with the continuing development and implementation of massively parallel sequencing technology. This sequencing revolution, however, has thus far had a relatively muted impact on studies of wild nonmodel organisms due largely to the difficulty of obtaining high-quality samples. This problem is particularly salient for endangered animals, cryptic animals, or animals for which it is otherwise difficult, undesirable, or unethical to obtain samples invasively.

Field researchers working with nonmodel animals have explored several noninvasive sample types for DNA analysis including feces, hair, urine, saliva, feathers, skin, and nails (Kohn and Wayne 1997). Of these, feces may be the most readily available in many taxa (Putman 1984). Indeed, since PCR amplification of DNA from feces was first demonstrated in the 1990s (Höss et al. 1992), noninvasive genetic studies from feces have revolutionized our understanding of the evolution, population structure, phylogeography, and behavior of nonmodel organisms. PCR amplification, however, is effective only for short sequences of DNA. The ability to generate cost-effective genomic-scale data of animals from feces using massively parallel sequencing would therefore constitute an important methodological advance towards bringing nonmodel organism studies into the genomic age.

Feces presents significant challenges for genetic analysis. DNA in feces is often fragmented and low in quantity. Fecal DNA extractions are further characterized by a frequent presence of co-extracted PCR inhibitors, sometimes complicating PCR detection of genotypes (Kohn and Wayne 1997), particularly with long amplicons. Finally, endogenous (host) DNA in feces constitutes a very low proportion, typically less than 5% (Qin et al. 2010; Perry et al. 2010; Snyder-Mackler et al. 2016), of total fecal DNA. Instead, fecal DNA contains a preponderance of DNA from exogenous (non-host) sources such as gut microbes, digesta, intestinal parasites, coprophagous animals, and other environmental organisms. Gut bacteria pose a particular challenge as they account for the highest proportion of DNA in feces (Perry et al. 2010; Qin et al. 2010).

Because of the high representation of exogenous DNA in feces, shotgun sequencing of fecal DNA would yield only a small proportion of reads matching the host genome. For genomic studies of host organisms, particularly those targeting populations, this represents a crippling obstacle in the presence of typical financial constraints. Without an effective enrichment procedure, sequencing of fecal DNA would be less efficient than that of invasively obtained “high-quality” DNA by at least one order of magnitude regardless of improvements in sequencing throughput or cost.

Attempts to enrich host DNA from feces for genomic analysis (Perry et al. 2010; Snyder-Mackler et al. 2016) have thus far employed targeted sequence capture methodologies. Sequence capture, like PCR, enriches DNA based on sequence specificity but unlike traditional PCR can work at any scale from a single locus (Whitney et al. 2004) to a whole genome (Melnikov et al. 2011; Carpenter et al. 2013; Snyder-Mackler et al. 2016). This method involves hybridizing DNA or RNA “baits,” either affixed to an array (Albert et al. 2007; Okou et al. 2007) or to magnetic beads in solution (Gnirke et al. 2009), to a mixture of target and nontarget sequences, thereby capturing targeted DNA from the mixture. Sequence capture has been used for instance to enrich human exomes (Ng et al. 2009), reduced-representation genomes (Suchan et al. 2016; Ali et al. 2016; Hoffberg et al. 2016), host DNA from ancient or museum specimens (Maricic et al. 2010; Carpenter et al. 2013; Mason et al. 2011; Bi et al. 2013), and pathogen genomes from human clinical samples (Melnikov et al. 2011). While the cost of custom oligonucleotide bait synthesis remains high, methods for transcribing custom baits from existing DNA templates (Melnikov et al. 2011; Carpenter et al. 2013) have driven costs significantly down, increasing sequence capture’s appeal.

Perry et al. (2010) first successfully enriched host DNA from feces at the genomic scale. Using a modified sequence capture employing custom-synthesized baits, they were able to highly enrich 1.5 megabases of chromosome 21, the X chromosome, and the mitochondrial genome from fecal samples of 6 captive chimpanzees. Their protocol, however, remains prohibitively expensive for population-level analysis due to the high cost of bait synthesis. More recently, Snyder-Mackler et al. (2016) performed whole-genome capture on fecal DNA, using RNA baits transcribed *in vitro* from high quality baboon samples to enrich host genomes from 62 wild baboons. Resulting libraries were sequenced to low coverage (mean 0.49×), but nevertheless provided sufficient information for reconstructing pedigree relationships.

Despite these methodological advances, targeted sequence capture has distinct drawbacks. To avoid the high cost of bait synthesis, RNA baits must first be transcribed from high-quality genomic DNA that is consumed by the process, limiting its appeal when working with species for which high-quality DNA is difficult to obtain or in short supply. The processes of both bait generation and hybridization with fecal DNA are labor-intensive and time-consuming, with the hybridization including an incubation step that alone takes 1 – 3 days (Snyder-Mackler et al. 2016). Because both RNA baits and the gDNA used to transcribe them are eventually depleted, the composition of RNA baits varies between bait sets, potentially impeding comparison of samples sequenced using different RNA baits and gDNA templates. *Trans* genomic captures (*i.e.* capturing DNA using baits from a different species) may complicate enrichment and introduce at least some capture biases (George et al. 2011), which will be a particular impediment for genomic studies for which high-quality DNA from related taxa is not accessible. Sequence capture may also introduce biases toward the capture of of low-complexity, highly repetitive genomic regions, as well as an excess of fragments from the mitochondrial genome (Samuels et al. 2013; Carpenter et al. 2013; Snyder-Mackler et al. 2016).

The present study exploits natural, evolutionarily ancient differences in CpG-methylation densities between vertebrate and bacterial genomes to enrich the host genome from feces, making noninvasive population genomics economically and practically feasible for the first time. This method, which we call FecalSeq, uses methyl-CpG-binding domain (MBD) proteins to selectively bind and isolate DNA with high CpG-methylation density. This enrichment method is inexpensive, *de novo*, and, crucially, captures target DNA without modification, thereby enabling downstream library preparation techniques including complexity reduction-based sequencing methods such as RADseq. Because of these properties, our method is well-suited for population genomic studies requiring high sequencing coverage, including those of nonmodel organisms for which few resources (e.g., high-quality samples or reference genomes) exist.

## RESULTS

Our method is a modification of a previously described technique for enriching the microbiome from vertebrate samples containing a majority of DNA from the host organism (Feehery et al. 2013). This technique employs a bait protein created by genetically fusing the human methyl-CpG binding domain protein 2 (MBD2) to the Fc tail of human IgG1. The resulting MBD2-Fc protein is then bound by a paramagnetic Protein A immunoprecipitation bead to create a complex that selectively binds double-stranded DNA with 5-methyl CpG dinucleotides. Because vertebrate DNA contains a high frequency of methylated CpGs (Hendrich and Tweedie 2003; Jabbari and Bernardi 2004) while bacterial DNA does not (Fang et al. 2012; Murray et al. 2012), this MBD bait complex selectively binds host DNA.

While Feehery et al. (2013) developed this method in order to remove contaminating host DNA for analysis of the microbiome, our strategy was to remove the contaminating fecal microbiome for analysis of host DNA. Therefore, after combining DNA with MBD baits, we retained the bound fraction with the goal of optimizing the selective recovery of host DNA (Fig. 1). Because our aim is to genotype populations with high coverage, we used the enriched host DNA to prepare double-digest RADseq libraries (Peterson et al. 2012) though with greater sequencing investment, the method in principle should work equally well for sequencing whole genomes.

To evaluate our approach, we enriched DNA extractions from the feces of 6 captive and 46 wild baboons, which we then used to prepare and sequence ddRADseq libraries. We also prepared ddRADseq libraries from blood-derived genomic DNA of all six captive baboons to facilitate controlled (same-individual) comparisons of blood and fecal libraries. All libraries were sequenced using Illumina sequencing.

Quantitative PCR estimates of starting host DNA proportions in fecal DNA extracts ranged widely, but were substantially lower in samples obtained from the wild (captive samples: mean 5.3%, range <0.01% – 17.4%; wild samples: mean 0.6%, range <0.01% – 4.9%; Supplemental Table S2).

**Fig. 1:**
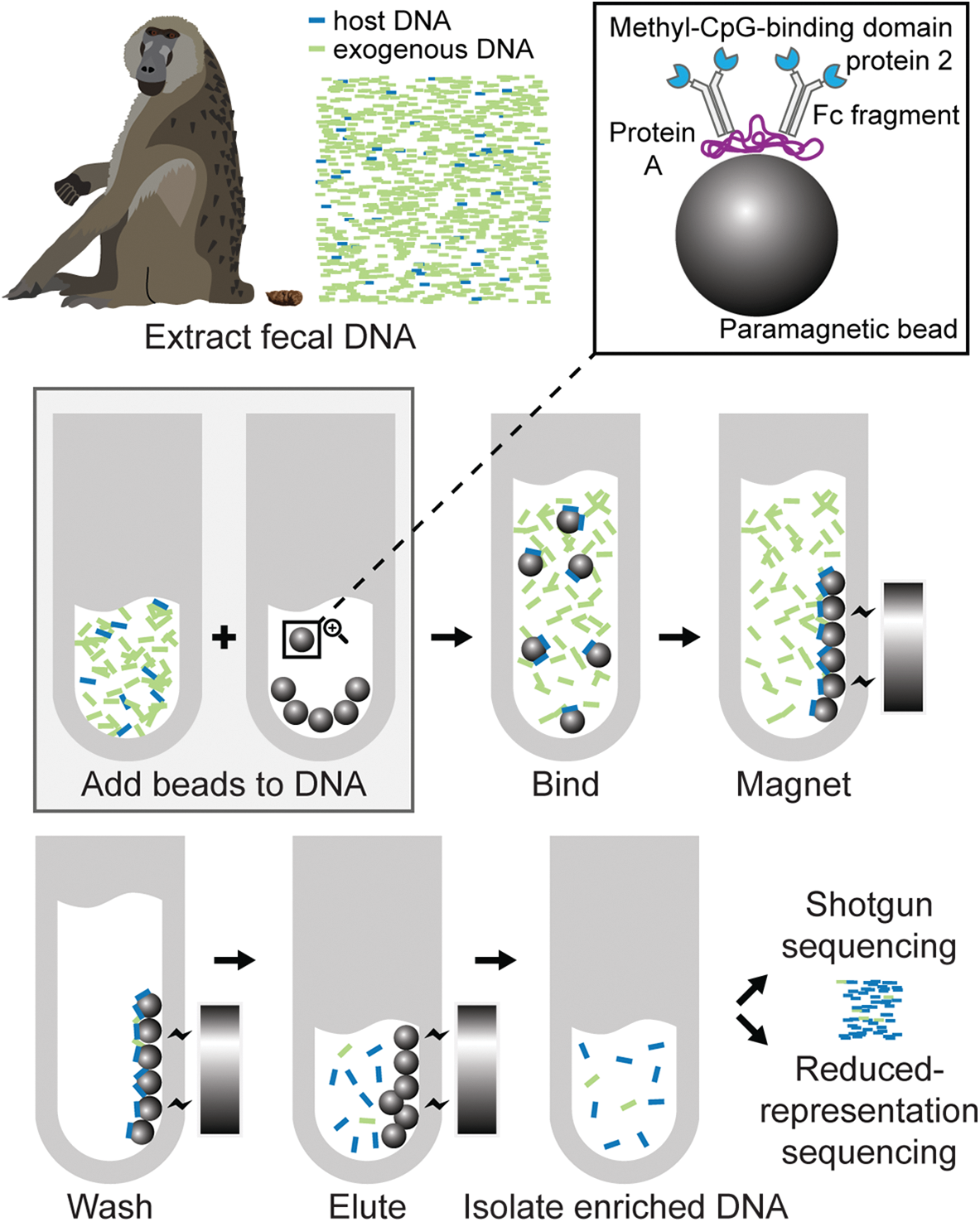
Overview of the FecalSeq method.

Based on two pilot libraries constructed from MBD-enriched fecal DNA, we found that there was large variation in the proportion of reads mapping to the baboon reference genome (mean 24.8%, range 0.7% – 81.2%; Supplemental Fig. S3; Supplemental Table S3), with the read mapping proportion correlating with starting host DNA proportions (library A: *r^2^* = 0.7338; *p* = 0.03; library B: *r^2^* = 0.9127, *p* < 0.01). Endogenous DNA proportions on average increased 13-fold (range 4.4 – 29.6; two samples removed due to starting proportions too low to quantify).

While some samples in our pilot libraries had high host DNA proportions following enrichment, these samples tended to already have high host DNA proportions prior to enrichment. Host DNA proportions following enrichment in the pilot libraries averaged only 4%, for instance, when samples with starting host DNA proportions greater than 1% were excluded. Because wild fecal DNA samples in our dataset on average started with less than 1% host DNA, we undertook a series of protocol optimization experiments to maximize the enrichment of these “low-quality” samples (Supplemental Table S5 and Supplemental Table S6).

Using a revised protocol based on our optimization experiments (Supplemental Protocol), we created and sequenced a third library from MBD-enriched fecal DNA. After noting substantial improvements in enrichment, we finally sequenced a fourth library with MBD-enriched fecal DNA from 40 wild baboons.

Despite having similar or even lower starting host DNA proportions, read mapping proportions in the third library were substantially higher than the prior two (mean 49.1%, range 8.9% – 75.3%; Fig. S3; Supplemental Table S3). Endogenous DNA proportions on average increased 318-fold (range 4.3 – 2632.2; one sample removed due to starting proportion too low to quantify).

The fourth library consisting entirely of fecal DNA from wild animals had the lowest starting concentrations of host DNA (mean 0.3%, range <0.01% – 3.1%). Following enrichment, however, host DNA proportions were nonetheless higher than our pilot libraries (mean 28.8%, range 1.5% – 73.6%; Supplemental Fig. S3; Supplemental Table S3). Endogenous DNA proportions on average increased 195-fold (range 23.7 – 486.9).

Overall, the revised protocol produced substantially higher enrichment, measured as fold increases in the proportion of host DNA, particularly for samples with very low starting proportions of host DNA (Fig. 2).While we sometimes were forced to use multiple rounds of extraction, thereby introducing variation in starting host proportions across same-individual trials, the revised protocol nonetheless exhibited robust improvement in read mapping proportions even when starting host proportions were substantially lower.

**Fig. 2:**
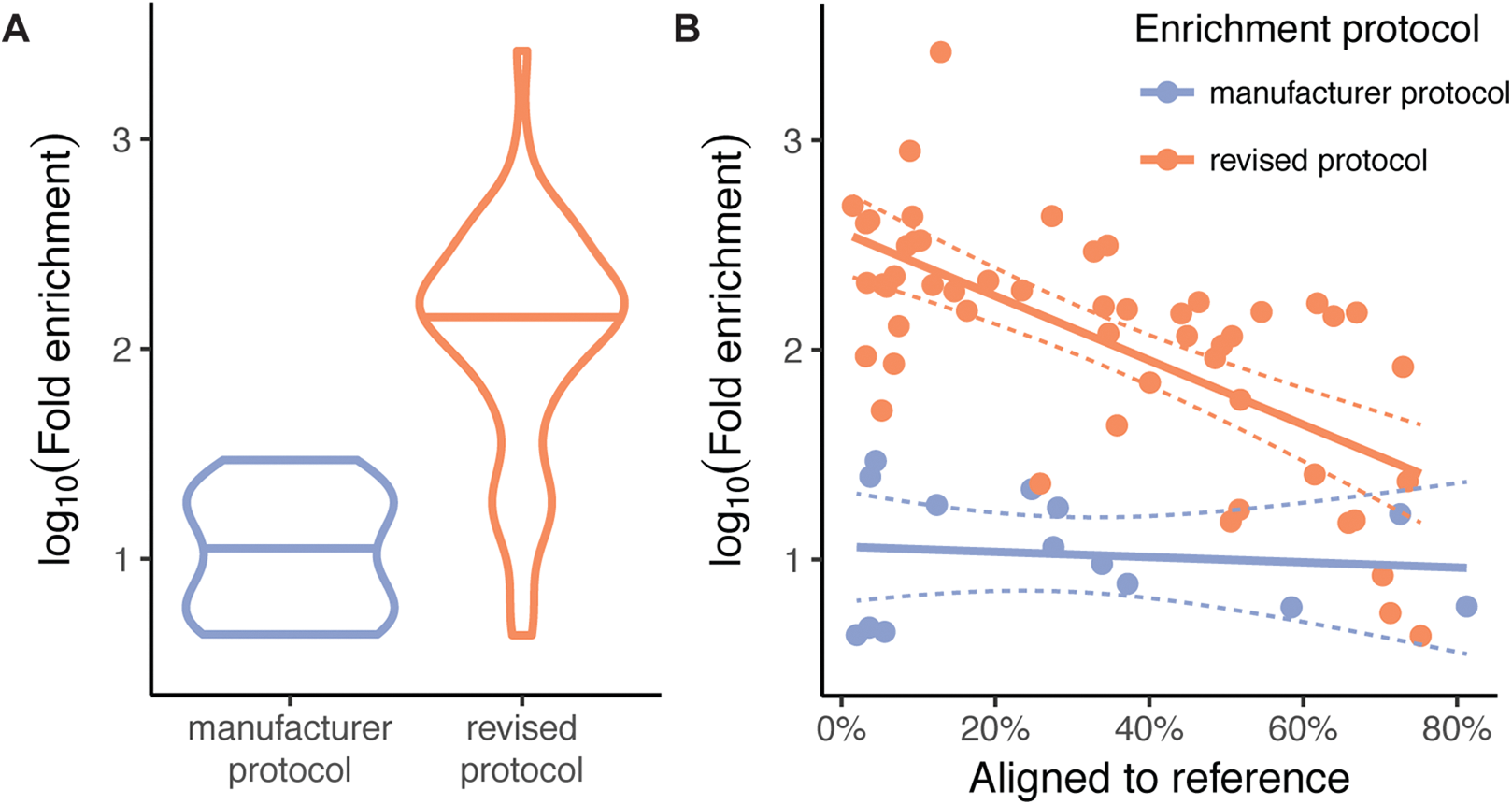
Comparison of the enrichment magnitude using the manufacturer protocol and the revised protocol. (a) Violin plots with the mean depicted show that the revised protocol results in substantially higher fold enrichment by approximately one order of magnitude. (b) A scatter plot shows that the revised protocol is particularly effective for samples with low starting quantities of host DNA. While some samples still had relatively small percentages of reads mapping to the baboon reference genome, these generally also exhibited the highest fold increases.

MBD binding may in principle select for genomic regions with relatively high CpG-methylation density, leading to dropout of other loci. Assessment of the concordance between blood- and feces-derived reads from the same individual was complicated by the correlation in ddRADseq between total reads and expected RADtags recovered and thereby SNPs discovered: a given RADtag is sequenced at a frequency inversely proportional to the deviation of its length from the mean of the size selection. Thus, we had to discern between dropout due to coverage-related stochasticity inherent in ddRADseq (Peterson et al. 2012) and that due to MBD enrichment. To perform this comparison, we computed the proportion of unique alleles between blood- and feces-derived RADtags from the same individual. For this test, we controlled for variation in sequencing coverage by randomly sampling reads as necessary in order to equalize total coverage among same-individual samples. Allelic dropout due to MBD enrichment would result in a higher proportion of alleles unique to blood-derived libraries relative to feces-derived libraries. We did not find a significant discrepancy (multi-sample-called SNPs: mean proportion unique alleles in blood = 2.3%, mean proportion unique alleles in feces = 2.3%; Wilcoxon signed rank test, *p* = 0.97; Fig. 3a).

**Fig. 3:**
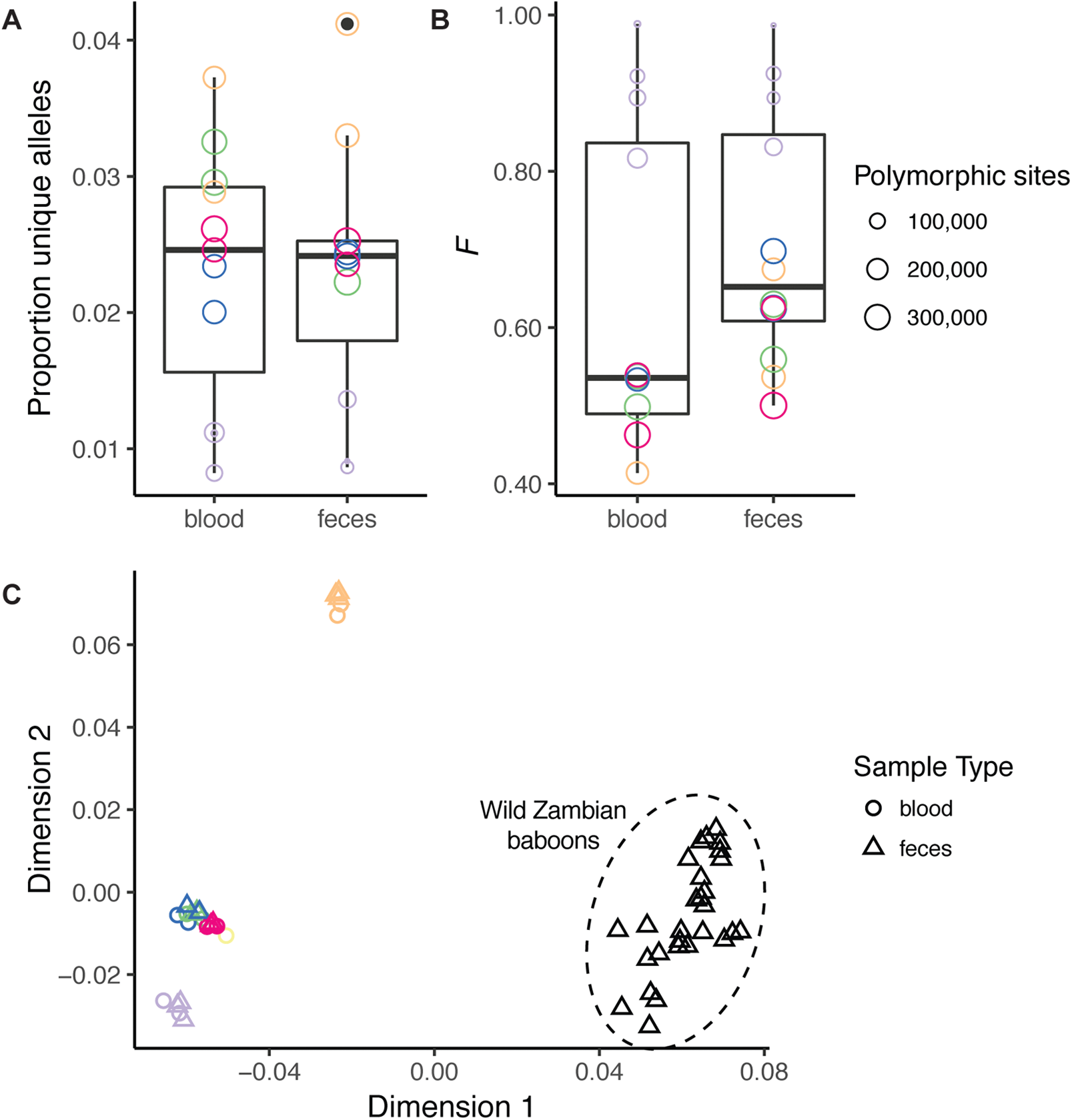
Concordance between blood- and feces-derived genotyping data from the same individuals. Colors symbolize the six captive individuals included in our study. Within these individuals, we did not find significant differences in (a) the proportion of unique alleles or (b) inbreeding coefficients from blood- and feces-derived libraries. The multidimensional scaling plot of identity-by-state shows (c) population structuring concordant with the known ancestry of animals (Supplemental Table S1). Distances between feces- and blood-derived sets of genotypes from the same individual are minimal, indicating that noise added by the enrichment method is dwarfed by the population structure signal in this baboon population dataset.

Dropout of entire RADtags is easily detectable given a reference genome or sufficient samples for comparison; dropout of a single allele at heterozygous sites is a more insidious potential bias. Allelic dropout due to MBD enrichment would result in a decrease in heterozygosity in MBD-enriched fecal libraries. Inbreeding coefficients (*F*) computed from same-individual RADtags exhibited in some cases higher values for feces-derived samples (Fig. 3b). This difference, however, was not statistically significant (mean *F*_blood_ = 0.63; mean *F*_feces_ = 0.71; Wilcoxon signed rank test, *p* = 0.47), indicating low allelic dropout attributable to the MBD enrichment. For this test, we also controlled for variation in sequencing coverage as described above.

As investigations of population structure are one potential application of our method, we visualized the wild and captive baboons’ identity-by-state via multidimensional scaling (MDS) using PLINK (Purcell et al. 2007; Chang et al. 2015), and confirmed that individuals clustered by their known species or ancestry and that blood- and feces-derived reads from the same individual were close together in the MDS space.

Stringent filtration of SNP sets, as would be implemented in a standard population genetic study, reduced the apparent biases attributable to fecal enrichment, measured both as total SNPs with a significant association with sample type (unfiltered: 25,079 out of 591,726, or 4.2%; filtered: 13 out of 7,202, or 0.2%) as well as total SNPs with significant missingness assessed via a chi-square test (unfiltered: 69,753 out of 550,224, or 12.7%; filtered: 0 out of 5,602, or 0%). Though more work is needed to quantify more exactly the extent and causal factors that lead to missingness, many population genetic analyses are robust to the low level of dropout our analyses reveal in addition to that which is inherent in the RADseq family of techniques (Gautier et al. 2013).

## DISCUSSION

Our methylation-based capture method achieves substantial enrichment of host DNA from fecal samples. Using our revised protocol developed through experimentation, we produced a mean 195-fold enrichment on our final library consisting entirely of fecal DNA obtained noninvasively under remote field conditions, with most samples nearly a decade old. A mean 28.8% of reads mapped to the baboon genome, despite starting with only a mean 0.34% of host DNA. Using fecal and blood DNA obtained from captive animals, we further demonstrate that feces-derived genotyping data following our method are concordant with corresponding data obtained from blood.

Feces are among the most readily accessible sources of information on wild animals (Kohn and Wayne 1997), and are particularly useful for population-level studies or studies of endangered or elusive species for which obtaining high-quality samples is difficult or undesirable. By exploiting methylation differences rather than sequence differences between host and bacterial DNA, FecalSeq is a *de novo* enrichment strategy that requires neither prior genome sequence knowledge nor the use of high-quality DNA for preparation of capture baits. This results in enrichment which is both inexpensive—we estimate a per-sample enrichment cost of $0.70 (Supplemental Note)—and replicable. The enrichment procedure is also relatively rapid and uncomplicated. Using a 96-well plate, we performed two sequential rounds of enrichment on all forty samples in our final library within a day (see Supplemental Protocol).

Importantly, FecalSeq is to our knowledge the first genomic-scale fecal DNA enrichment method that is compatible with most downstream library preparation methods for massively parallel sequencing. Through our use of ddRADseq, we demonstrate that our method facilitates low-cost high-capacity genotyping of wild populations without introducing significant bias. Further, because ddRADseq is customizable (Peterson et al. 2012), there is substantial flexibility for researchers to optimize the number of samples and the fraction of the genome sequenced for particular research questions. This is not possible for libraries prepared using targeted sequence capture, which are therefore currently limited mainly to low-coverage analyses at the population level (Snyder-Mackler et al. 2016). Transcription of sequence capture baits from reduced-representation libraries may potentially help address this problem (Suchan et al. 2016; Ali et al. 2016; Hoffberg et al. 2016), but its efficacy for fecal DNA has yet to be demonstrated.

Double-digest RADseq is possible for genotyping species with or without a reference genome. We aligned our sequencing reads to the baboon reference genome for this study, but our approach is likely also applicable to species without a reference genome. In these cases, an additional pre-screening step would be necessary, in which exogenous reads are filtered out through comparison to the nearest available genome, before proceeding to clustering and variant identification as per normal reference-free ddRAD methods

We robustly found that sequencing efficiency (percentage of reads assigned to target genome) of MBD-enriched fecal DNA libraries correlates strongly with starting proportions of host DNA, echoing findings using other capture methods (Snyder-Mackler et al. 2016). Future attention should therefore be directed towards fecal sample collection, storage, and extraction methods that maximize the selective recovery of host nuclear DNA (e.g., Ramón-Laca et al. 2015). While we demonstrated effective genotyping of samples with often very low starting proportions of host DNA (the vast majority < 0.5%), future studies may consider pre-screening extracted DNA samples using qPCR to select for samples with high starting proportions of host DNA.

Low starting proportions of host DNA present a challenge not only because they result in lower sequencing efficiency, but also because they correlate with low absolute quantities of DNA belonging to the host organism. In some cases, particularly in samples collected from wild animals under field conditions, starting proportions of host DNA were so low that only approximately 0.1 ng of target DNA was available in a 1 µg fecal DNA extract. Given the large genome sizes of baboons (approximately 3 Gb) and many other vertebrates, substantial allelic dropout is expected in these cases. Significantly, this challenge cannot be fully addressed by this or any other enrichment method and remains an important consideration for researchers working with feces. It can be minimized, however, by optimizing the enrichment procedures to maximize the recovery of target DNA present in a fecal DNA sample, as well as by increasing the total amount of starting fecal DNA.

Because MBD enrichment partitions DNA based on CpG-methylation density, FecalSeq does not enrich hypomethylated host mitochondrial DNA (Rebelo et al. 2009). While this may be undesirable for studies requiring the matrilineally inherited marker, it also precludes the disproportionately high representation of mitochondrial DNA that is typical in libraries prepared using the targeted sequence capture approach (Perry et al. 2010; Samuels et al. 2013; Carpenter et al. 2013; Snyder-Mackler et al. 2016). FecalSeq may, however, co-enrich nuclear DNA from exogenous eukaryotes such as from plant or animal digesta. Care should therefore be taken to minimize the presence of exogenous eukaryotic tissues or cells, although the degree to which this is a problem in practice is currently unknown. As cell-wall-bound plant cells may be more likely to pass through the digestive tract intact, extraction methods that minimize lysis of cell walls should be preferred. We speculate that prey DNA from carnivorous animals may be more difficult to partition from host DNA.

Since PCR amplification of DNA from feces was first achieved in the 1990s (Höss et al. 1992; Constable et al. 1995; Gerloff et al. 1995), noninvasive genetic studies have revolutionized our understanding of the evolution, ecology, and behavior of nonmodel organisms. By facilitating low-cost genomic-scale sequencing from feces, our method connects a community of field researchers with the benefits of massively parallel sequencing, ushering noninvasive organism studies into the genomic age.

## METHODS

### Samples

Blood and fecal samples were collected from six captive baboons (genus *Papio*) housed at the Southwest National Primate Research Center (SNPRC) at the Texas Biomedical Research Institute. The individuals were of either *P. anubis* or hybrid ancestry (Supplemental Table S1). All six baboons were fed a diet manufactured by Purina LabDiet (“Monkey Diet 15%”) containing 15% minimum crude protein, 4% minimum crude fat, and 10% maximum crude fiber. In separate sedation events, blood and feces were collected from the same individual who was isolated for the duration of the sedation. Following centrifugation, the buffy coat was isolated from blood samples and stored at -80 °C. 2 ml of feces were also collected into 8 ml tubes containing 4 ml of RNALater (Ambion). All procedures were conducted under the Texas Biomedical Research Institute IACUC protocol #1403 PC 0. Sedation and blood draws were performed under the supervision of a veterinarian and animals were returned immediately to their enclosures following recovery.

In addition, we collected or obtained fecal samples from 46 wild baboons in Zambia. Samples were collected between 2006 and 2015 from the Luangwa Valley, the Lower Zambezi National Park, Choma, or Kafue National Park and are of *P. kindae × P. cynocephalus*, *P. griseipes*, or *P. kindae × P. griseipes* ancestry (Supplemental Table S1). As with the SNPRC samples, 2 ml of feces were collected into 8 ml tubes containing 4 ml of RNALater. In contrast to the SNPRC samples, however, these samples were collected noninvasively from unhabituated animals in remote field conditions. Samples therefore could not be attributed to particular animals, although samples were selected to avoid duplication using either field observations or geographic distance. Following collection, samples were stored without refrigeration for 1 – 6 months before being frozen at -20 °C for long-term storage.

Buffy coat extractions were performed using the QIAamp DNA Blood Mini Kit (Qiagen), following manufacturer’s instructions. Fecal extractions were performed using the QIAamp DNA Stool Mini Kit (Qiagen) following manufacturer’s instructions for optimizing host DNA yields. DNA concentration and yield were measured using a Qubit dsDNA BR Assay (Life Technologies). In some cases, multiple DNA extractions from the same individuals were necessary when DNA was depleted over the course of this study.

We estimated the proportion of host DNA for each fecal DNA extraction using quantitative PCR (qPCR) by comparing estimates of host DNA concentration obtained by qPCR to estimates of total fecal DNA concentration obtained by Qubit. Amplification was conducted using universal mammalian *MYCBP* primers (Morin et al. 2001) and evaluated against a standard curve constructed from the liver DNA of an individual baboon. Samples and standards were run in duplicate alongside positive and negative controls (see Supplemental Protocol for full details).

### DNA enrichment

DNA was enriched using the NEBNext Microbiome DNA Enrichment Kit (New England BioLabs). This enrichment procedure (Feehery et al. 2013) captures eukaryotic DNA using a methylated CpG-specific binding domain protein fused to the Fc fragment of human IgG (MBD2-Fc) to selectively target sequences with high CpG methylation density.

MBD2-Fc-bound magnetic beads were prepared according to manufacturer instructions in batches ranging from 40 to 160 µl. For each *n* µl batch, we prebound 0.1 *× n* µl MBD2-Fc protein to *n* µl protein A magnetic beads by incubating the mixture with rotation for 10 min at room temperature. The bound MBD2-Fc magnetic beads were then collected by magnet and washed twice with 1 ml ice-cold 1X bind/wash buffer before being resuspended in *n* µl ice-cold 1X bind/wash buffer.

As a pilot experiment, we prepared two successive libraries, library A and library B, following manufacturer’s instructions for capturing methylated host DNA, with minor protocol modifications incorporated for the second pilot library (library B). Library A included MBD-enriched fecal DNA from 4 SNPRC baboons and 2 Luangwa Valley baboons, as well as blood DNA from the same SNPRC baboons. Library B included MBD-enriched fecal DNA from 4 SNPRC baboons (with two repeats from library A), 4 Kafue National Park baboons, and 2 Luangwa Valley baboons, as well as blood DNA from 2 SNPRC baboons. For each fecal DNA sample, we combined 1 – 2 µg of extracted fecal DNA with 160 µl of prepared protein-bound beads and a variable volume of ice-cold 5X bind/wash buffer for maintaining 1X concentration of bind/wash buffer. After combining beads and DNA, we incubated the mixture at room temperature with rotation for 15 min. DNA and MBD2-Fc-bound magnetic beads were then collected by magnet and the supernatant removed. For samples in library A, we washed the collected beads with 1 ml of ice-cold 1X bind/wash buffer. For samples in library B, we conducted three expanded wash steps to maximize the removal of unbound DNA. For each wash in library B, we added 1 ml of ice-cold 1X bind/wash buffer and mixed the beads on a rotating mixer for three minutes at room temperature before collecting the beads by magnet and removing the supernatant. Following the final wash, we resuspended and incubated the beads at 65 °C with 150 µL of 1X TE buffer and 15 µL of Proteinase K for 20 min with occasional mixing. The eluted DNA was then separated by magnet, purified with 1.5X homemade SPRI bead mixture (Rohland and Reich 2012), and quantified using a Qubit dsDNA HS Assay (Life Technologies).

Our pilot sequencing results from libraries A and B revealed large variation in the percentage of reads mapping to the baboon genome, with mapping percentages ranging from 1.1% to 79.3%, with much of the variation correlating with the proportion of host DNA in the unenriched fecal DNA sample (Supplemental Fig. S1). To expand the utility of the enrichment protocol to all fecal DNA samples, we conducted a series of capture experiments designed to optimize the enrichment of host DNA from “low-quality” samples (i.e., samples with low proportions of host DNA). For these experiments, we artificially simulated fecal DNA by combining high-quality baboon liver or blood genomic DNA (liver: SNPRC ID #19334; blood: SNPRC ID #14068 or #25567) with *E. coli* DNA (K12 or ATCC 11303 strains) at controlled proportions. The resulting post-enrichment proportion of baboon and *E. coli* DNA was evaluated by qPCR in two analyses using (1) universal mammalian *MYCBP* (Morin et al. 2001) and (2) universal bacterial 16S rRNA (16S) (Corless et al. 2000) primers along with standards created from the same respective organisms (experiments and results are described in detail in Supplemental Table S2).

We prepared a third and fourth library, libraries C and D, incorporating modifications (Supplemental Protocol) based on results from our capture optimization experiments. For these captures, we added a much smaller volume of prepared MBD2-Fc-bound magnetic beads (1 – 22 µl) based on the estimated proportion of starting host DNA, kept the capture reaction volume consistent at a relatively low 40 µl (concentrating samples as needed using a SPRI bead cleanup), added an extra wash step in which samples were resuspended in 100 µl of 1X bind/wash buffer then incubated at room temperature for 3 minutes with rotation, and eluted samples in 100 µl 2 M NaCl. For four fecal DNA samples in library C and all of library D, we serially enriched the samples by repeating the capture reaction with 30 µl of MBD-enriched DNA (post SPRI-bead cleanup). Library C included fecal DNA from 5 SNPRC baboons, 2 Kafue National Park baboons, and 1 Luangwa Valley baboon. Library D contained fecal DNA from 6 Lower Zambezi National Park baboons, 4 Choma baboons, and 30 Kafue National Park baboons.

We prepared a final library, library E, from independently extracted blood DNA from five SNPRC baboons in order to quantify the stochasticity associated with independent library preparation from independent extracts. The composition of libraries A-E are described in detail in Supplemental Table S2 and Supplemental Table S3.

### Library preparation and sequencing

Library preparation followed standard double-digest restriction site-associated DNA sequencing (ddRAD-seq) procedures (Peterson et al. 2012) with modifications to accommodate low input as described below.

For all samples, including blood DNA and MBD-enriched fecal DNA, we digested DNA with *Sph*I and *Mlu*CI (New England Biolabs), following a ratio of 1 unit of each enzyme per 20 ng of DNA. Enzymes were diluted up to 10X using compatible diluents (New England Biolabs) to facilitate pipetting of small quantities, using an excess of enzyme if necessary to avoid pipetting less than 1 µl of the diluted enzyme mix. As the total amount of post-enrichment fecal DNA is by nature low, we adjusted adapter concentrations in the ligation reaction to ~0.1 µM for barcoded P1 and ~3 µM for P2, which correspond to excesses of adapters between 1 – 2 orders of magnitude. Since adapter-ligated samples are multiplexed into pools in equimolar amounts, we made efforts to combine samples with similar concentrations and enrichment when known. We used the BluePippin (Sage Science) with a 1.5% agarose gel cassette for automated size selection of pooled individuals, with a target of 300 bp (including adapters) and extraction of a “tight” collection range. For PCR amplification, we ran all reactions in quadruplicate to minimize PCR biases and attempted to limit the number of PCR cycles. As the concentration of post-size-selection pools was below the limits of detection without loss of a considerable fraction of the sample, estimation of the required number of PCR cycles was difficult. We therefore iteratively quantified products post-PCR and added cycles as necessary. The total number of PCR cycles per pool is reported in Supplemental Table S3, but was usually 24. Finally, libraries were sequenced using either Illumina MiSeq (libraries A-C; 2 × 150 paired-end) or Illumina HiSeq 2500 (library D; 2 × 100 paired-end) sequencing with 30% spike in of PhiX control DNA.

### Analysis

We demultiplexed reads by sample and mapped them to the baboon reference genome (papAnu2; Baylor College of Medicine Human Genome Sequencing Center) using BWA with default parameters and the BWA-ALN algorithm (Li and Durbin 2009). For every pair of blood and fecal samples from the same individual, we down-sampled mapped reads to create new pairs with equal coverage in order to control for biases due to differences in sequencing depth. After realignment around indels, we identified variants using GATK UnifiedGenotyper (DePristo et al. 2011), in parallel analyses (1) calling variants in all samples at once and (2) processing each sample in isolation to avoid biasing variant calls from other samples at the expense of accuracy. Homozygous sites matching the reference genome were listed as missing when variants were inferred in single individuals. Variants were filtered with GATK VariantFiltration (filters: `QD < 2.0, MQ < 40.0, FS > 60.0, HaplotypeScore > 13.0, MQRankSum <   −12.5, ReadPosRankSum < −8.0`) and indels were excluded.

We digested the baboon reference genome *in silico*, tallied reads within each predicted RADtag, and gathered the following information about each region: length, GC percentage, and CpG count in region ± 5 kb. We also calculated read depth in these simulated RADtags. Distributions of blood and fecal RADtags’ length, GC percentage, and local CpG density (Supplemental Fig. S2) were visually inspected for gross distortion due to widespread dropout.

If the fecal enrichment procedure caused widespread allelic dropout, the proportion of alleles unique to the blood samples would be higher than that to the fecal sample. We tallied these unique alleles with VCFtools (Danecek et al. 2011) and tested for an excess with a Wilcoxon signed rank test.

To quantify loss of heterozygosity due to allelic dropout, we computed the inbreeding coefficient, *F* for all blood-feces pairs with equalized coverage, using both the individually called and multi-sample called SNP sets. The presence of dropout is expected to inflate *F*. We tested for differences in paired samples’ estimates of *F* via a Wilcoxon signed rank test. The dataset is not filtered for missingness, so sequencing errors inferred to be true variants may inflate heterozygosity estimates, thus deflating *F*.

To create a stringently filtered dataset with high genotyping rate, we filtered the multi-sample called SNPs in PLINK (Purcell et al. 2007; Chang et al. 2015), retaining only those genotyped in at least 90% of samples and removing samples with genotypes at fewer than 10% of sites. This filtered set was further pruned for linkage disequilibrium by sliding a window of 50 SNPs across the chromosome and removing one random SNP in each pair with *r^2^* > 0.5. Using all samples, we performed multidimensional scaling to visualize identity by state (IBS). Using just the samples that were part of the same-individual blood-feces pairs, we then performed an association test and missingness chi-square test to detect allele frequencies or missingness that correlated with sample type. We did the same with the unfiltered dataset as well. Though we had few pairs of fecal samples from the same individual, we computed distance between pairs of samples from the same individual using the stringently filtered dataset to compare distance between and within sample types via a Wilcoxon signed rank test.

## ACKNOWLEDGMENTS

We thank Clifford Jolly and Jane Phillips-Conroy for contributing fecal samples collected in 2006 – 2007 from Zambia. We also thank the Zambia Wildlife Authority (now the Department of National Parks & Wildlife) and the University of Zambia for granting permission and providing support for fieldwork. We thank Erbay Yigit, Andrew Burrell, and Todd Disotell for helpful conversations. This study was funded by the National Science Foundation (BCS 1341018, BCS 1260816, BCS 1029302, SMA 1338524), the Leakey Foundation, the Wenner-Gren Foundation, the National Geographic Society, and the NYU University Research Challenge Fund. The Genome Technology Center at NYU is supported by NIH/NCATS UL1 TR00038 and NIH/NCI P30 CA016087. K.L.C. is supported by NSF fellowship DGE 1143954.

## AUTHOR CONTRIBUTIONS

K.L.C. and C.M.B. conceived the project and wrote the paper. K.L.C. collected samples and led the labwork. C.M.B. led the analysis.

---

**Supplemental Fig. S1:**
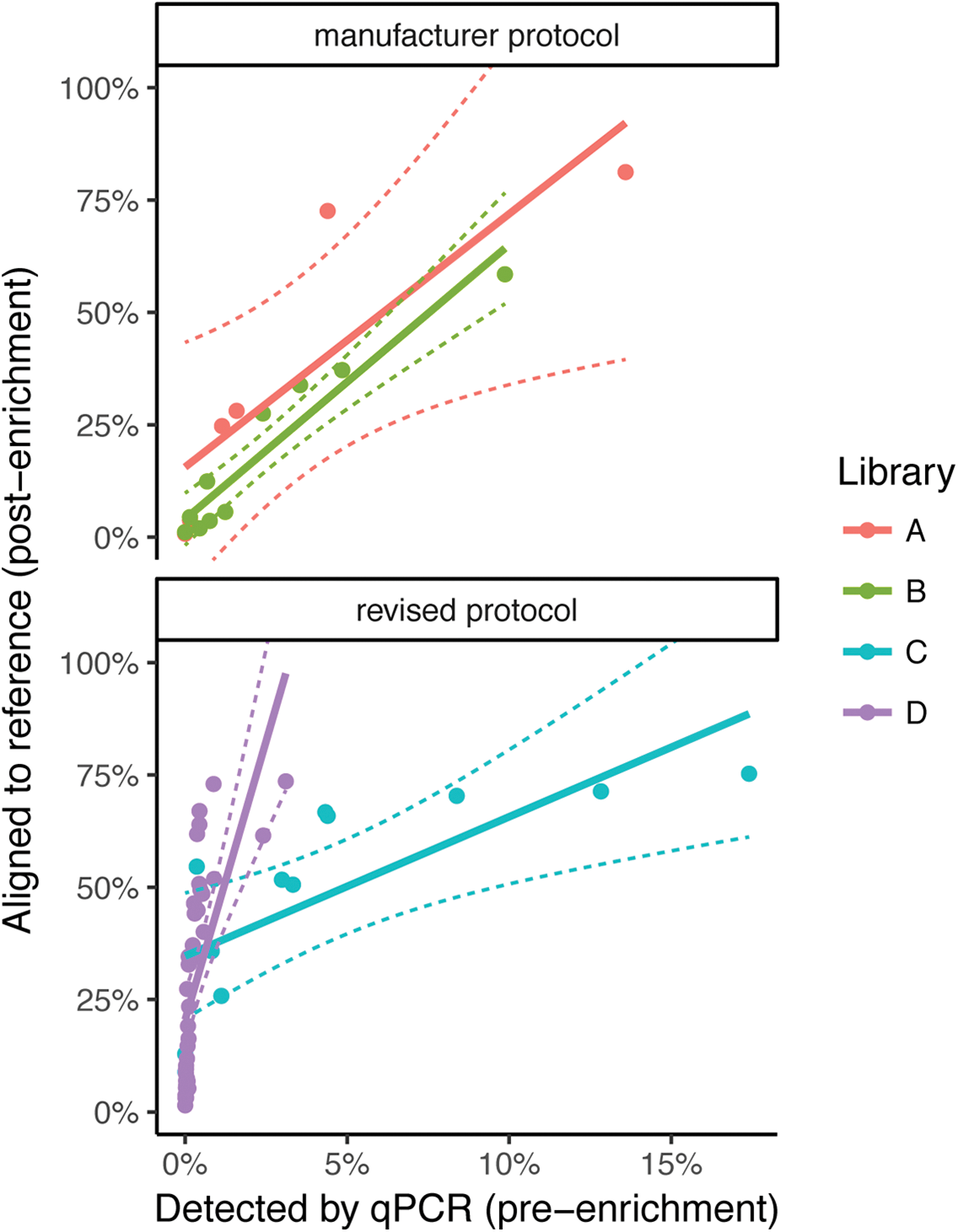
Relationship between pre-enrichment host percentage, as estimated by quantitative PCR, and post-enrichment host percentage, as estimated by alignment of sequencing reads to the baboon reference genome.

**Supplemental Fig. S2:**
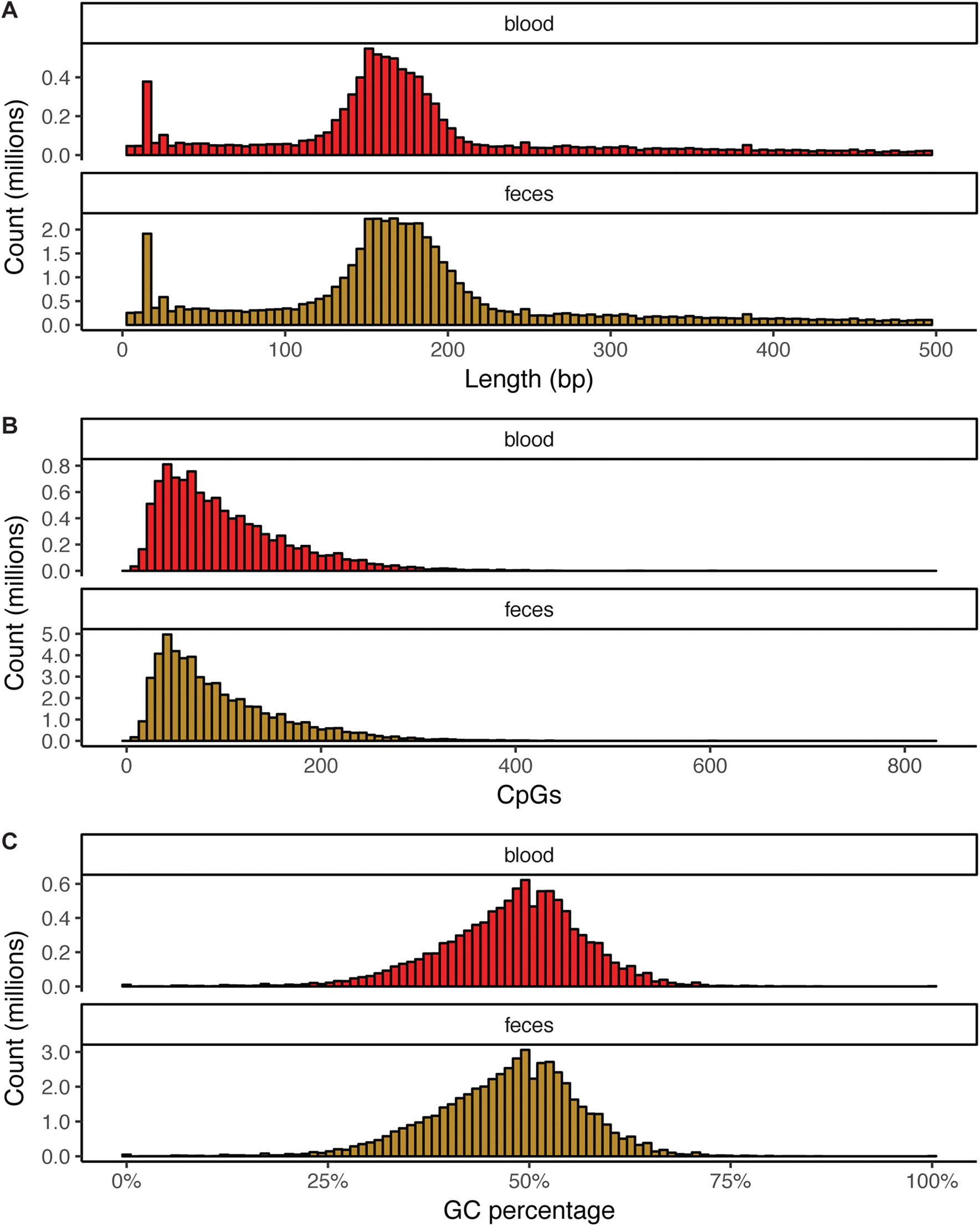
Combined distributions of (a) RADtag lengths, (b) CpG counts (within the boundaries of the sequenced RADtag ±5, 000), and (c) GC percentages in sequenced libraries.

**Supplemental Fig. S3:**
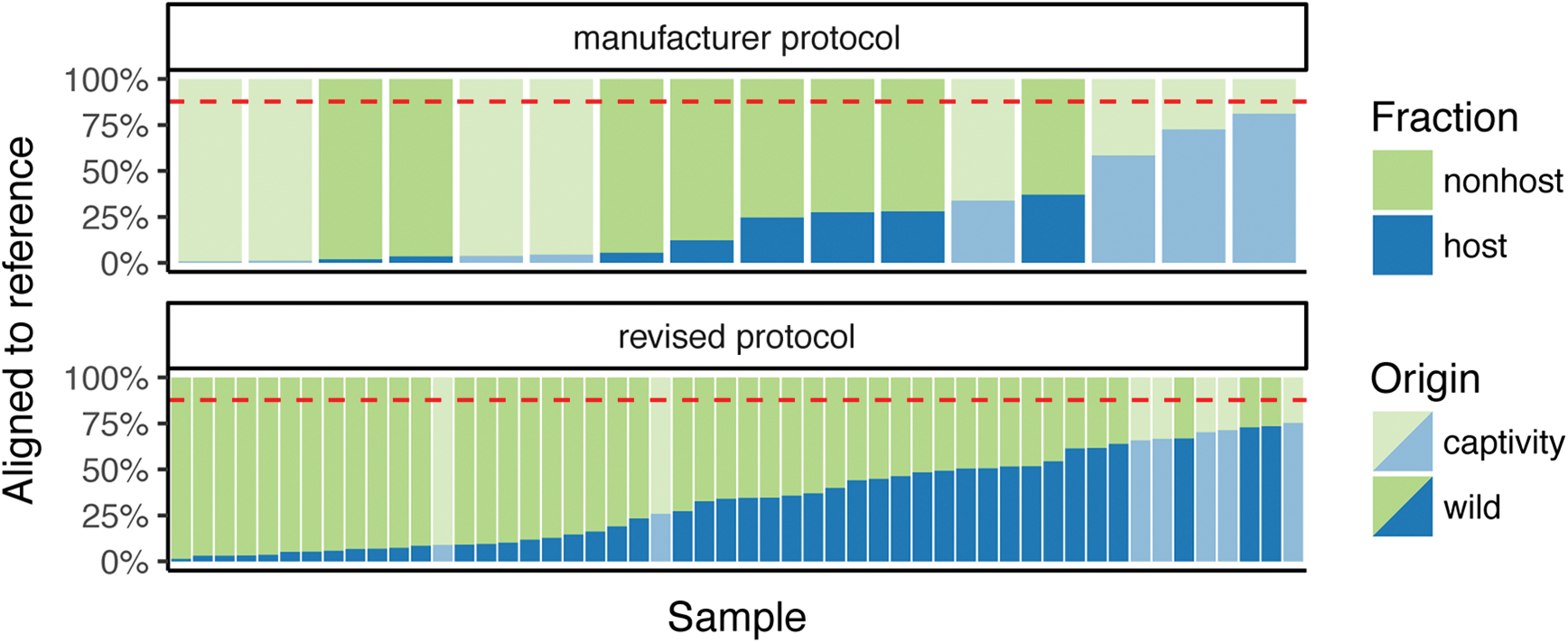
Percentage of reads mapping to the baboon reference genome (papAnu2) for all samples included in this study. Sixteen samples were enriched using the manufacturer protocol and 52 using the revised protocol.

**Supplemental Table S1:** Animals sequenced for this study.

| Individual | Locale | Taxon | Sex | Origin | Year |
| --- | --- | --- | --- | --- | --- |
| SNPRC #13245 | SNPRC | *Papio anubis* | male | captivity | 2014 |
| SNPRC #14068 | SNPRC | *Papio anubis* | male | captivity | 2014 |
| SNPRC #25567 | SNPRC | *Papio anubis* × *Papio ursinus* | female | captivity | 2014 |
| SNPRC #27278 | SNPRC | *Papio anubis* × *Papio cynocephalus* | male | captivity | 2014 |
| SNPRC #27958 | SNPRC | *Papio anubis* | female | captivity | 2014 |
| SNPRC #28064 | SNPRC | *Papio anubis* | female | captivity | 2014 |
| BZ06-051 | South Luangwa NP | *Papio kindae* × *Papio cynocephalus* | unknown | wild | 2006 |
| BZ06-053 | South Luangwa NP | *Papio kindae* × *Papio cynocephalus* | unknown | wild | 2006 |
| BZ06-066 | South Luangwa NP | *Papio kindae* × *Papio cynocephalus* | unknown | wild | 2006 |
| BZ06-148 | North Luangwa NP | *Papio kindae* × *Papio cynocephalus* | unknown | wild | 2006 |
| BZ06-218 | Lower Zambezi NP | *Papio griseipes* | unknown | wild | 2006 |
| BZ06-220 | Lower Zambezi NP | *Papio griseipes* | unknown | wild | 2006 |
| BZ06-221 | Lower Zambezi NP | *Papio griseipes* | unknown | wild | 2006 |
| BZ06-224 | Lower Zambezi NP | *Papio griseipes* | unknown | wild | 2006 |
| BZ06-225 | Lower Zambezi NP | *Papio griseipes* | unknown | wild | 2006 |
| BZ06-227 | Lower Zambezi NP | *Papio griseipes* | unknown | wild | 2006 |
| BZ07-001 | Choma | *Papio griseipes* | unknown | wild | 2007 |
| BZ07-004 | Choma | *Papio griseipes* | unknown | wild | 2007 |
| BZ07-005 | Choma | *Papio griseipes* | unknown | wild | 2007 |
| BZ07-007 | Choma | *Papio griseipes* | unknown | wild | 2007 |
| BZ07-029 | Kafue NP | *Papio kindae* × *Papio griseipes* | unknown | wild | 2007 |
| BZ07-030 | Kafue NP | *Papio kindae* × *Papio griseipes* | unknown | wild | 2007 |
| BZ07-032 | Kafue NP | *Papio kindae* × *Papio griseipes* | unknown | wild | 2007 |
| BZ07-034 | Kafue NP | *Papio kindae* × *Papio griseipes* | unknown | wild | 2007 |
| BZ07-035 | Kafue NP | *Papio kindae* × *Papio griseipes* | unknown | wild | 2007 |
| BZ07-039 | Kafue NP | *Papio kindae* × *Papio griseipes* | unknown | wild | 2007 |
| BZ07-041 | Kafue NP | *Papio kindae* × *Papio griseipes* | unknown | wild | 2007 |
| BZ07-042 | Kafue NP | *Papio kindae* × *Papio griseipes* | unknown | wild | 2007 |
| BZ07-045 | Kafue NP | *Papio kindae* × *Papio griseipes* | unknown | wild | 2007 |
| BZ07-047 | Kafue NP | *Papio kindae* × *Papio griseipes* | unknown | wild | 2007 |
| BZ07-100 | Kafue NP | *Papio kindae* × *Papio griseipes* | unknown | wild | 2007 |
| Chiou-14-001 | Kafue NP | *Papio kindae* × *Papio griseipes* | unknown | wild | 2014 |
| Chiou-14-003 | Kafue NP | *Papio kindae* × *Papio griseipes* | unknown | wild | 2014 |
| Chiou-14-004 | Kafue NP | *Papio kindae* × *Papio griseipes* | unknown | wild | 2014 |
| Chiou-14-005 | Kafue NP | *Papio kindae* × *Papio griseipes* | unknown | wild | 2014 |
| Chiou-14-030 | Kafue NP | *Papio kindae* × *Papio griseipes* | unknown | wild | 2014 |
| Chiou-14-036 | Kafue NP | *Papio kindae* × *Papio griseipes* | unknown | wild | 2014 |
| Chiou-14-039 | Kafue NP | *Papio kindae* × *Papio griseipes* | unknown | wild | 2014 |
| Chiou-14-041 | Kafue NP | *Papio kindae* × *Papio griseipes* | unknown | wild | 2014 |
| Chiou-14-042 | Kafue NP | *Papio kindae* × *Papio griseipes* | unknown | wild | 2014 |
| Chiou-14-044 | Kafue NP | *Papio kindae* × *Papio griseipes* | unknown | wild | 2014 |
| Chiou-14-050 | Kafue NP | *Papio kindae* × *Papio griseipes* | unknown | wild | 2014 |
| Chiou-14-054 | Kafue NP | *Papio kindae* × *Papio griseipes* | unknown | wild | 2014 |
| Chiou-14-056 | Kafue NP | *Papio kindae* × *Papio griseipes* | unknown | wild | 2014 |
| Chiou-14-057 | Kafue NP | *Papio kindae* × *Papio griseipes* | unknown | wild | 2014 |
| Chiou-14-058 | Kafue NP | *Papio kindae* × *Papio griseipes* | unknown | wild | 2014 |
| Chiou-14-059 | Kafue NP | *Papio kindae* × *Papio griseipes* | unknown | wild | 2014 |
| Chiou-14-065 | Kafue NP | *Papio kindae* × *Papio griseipes* | unknown | wild | 2014 |
| Chiou-14-069 | Kafue NP | *Papio kindae* × *Papio griseipes* | unknown | wild | 2014 |
| Chiou-15-003 | Kafue NP | *Papio kindae* × *Papio griseipes* | unknown | wild | 2015 |
| Chiou-15-004 | Kafue NP | *Papio kindae* × *Papio griseipes* | unknown | wild | 2015 |
| Chiou-15-005 | Kafue NP | *Papio kindae* × *Papio griseipes* | unknown | wild | 2015 |

**Supplemental Table S2:** Fecal DNA enrichment results. Key: ID, capture experiment ID; Lib, library ID; Ind, individual (see Supplemental Table S1); PHB, percent host DNA before; TD, total fecal DNA used (ng); BV, bead volume used (µl); TV, total reaction volume (µl); TY, total DNA yield (ng); NE, number of enrichment steps; PHA, percent host DNA after; NDB, *n*-fold decrease in bacterial DNA.

| ID | Lib | Ind | PHB | TD | BV | TV | TY | NE | PHA | NDB |
| --- | --- | --- | --- | --- | --- | --- | --- | --- | --- | --- |
| A01F.T002 | A | SNPRC #14068 | 0.15% | 1,000.00 | 160.00 | 163.00 | 82.83 | single | 6.79% | - |
| A02F.T005 | A | SNPRC #25567 | 4.40% | 2,000.00 | 160.00 | 174.80 | 455.40 | single | 50.90% | - |
| A03F.T007 | A | SNPRC #27278 | 0.00% | 1,000.00 | 160.00 | 165.10 | 191.40 | single | 6.38% | - |
| A04F.T009 | A | SNPRC #27958 | 13.59% | 2,000.00 | 160.00 | 172.40 | 584.10 | single | 47.50% | - |
| A05F.B051 | A | BZ06-051 | 1.59% | 1,800.00 | 160.00 | 285.00 | 122.43 | single | - | - |
| A06F.B053 | A | BZ06-053 | 1.14% | 1,365.00 | 160.00 | 535.00 | 119.79 | single | - | - |
| B01F.T001 | B | SNPRC #13245 | 3.55% | 1,000.00 | 160.00 | 170.90 | 72.80 | single | 19.45% | - |
| B02F.T002 | B | SNPRC #14068 | 0.15% | 1,000.00 | 160.00 | 164.40 | 16.08 | single | 8.34% | - |
| B03F.T007 | B | SNPRC #27278 | 0.00% | 1,000.00 | 160.00 | 165.40 | 26.80 | single | 12.68% | - |
| B04F.T010 | B | SNPRC #28064 | 9.87% | 1,000.00 | 160.00 | 177.50 | 78.80 | single | 125.38% | - |
| B05F.C030 | B | Chiou-14-030 | 1.24% | 1,000.00 | 160.00 | 285.00 | 38.16 | single | 14.31% | - |
| B06F.C050 | B | Chiou-14-050 | 2.40% | 1,000.00 | 160.00 | 198.10 | 13.28 | single | 10.81% | - |
| B07F.C065 | B | Chiou-14-065 | 0.76% | 1,000.00 | 160.00 | 285.00 | 22.96 | single | 25.44% | - |
| B08F.C069 | B | Chiou-14-069 | 0.45% | 1,000.00 | 160.00 | 211.20 | 15.92 | single | 17.88% | - |
| B09F.B066 | B | BZ06-066 | 4.85% | 1,000.00 | 160.00 | 285.00 | 39.20 | single | 38.72% | - |
| B10F.B148 | B | BZ06-148 | 0.68% | 1,000.00 | 160.00 | 292.10 | 9.92 | single | 9.07% | - |
| C01F.T001 | C | SNPRC #13245 | 4.33% | 1,000.00 | 6.92 | 40.00 | 20.80 | single | 65.00% | 1.89 |
| C02F.T002 | C | SNPRC #14068 | 1.12% | 1,000.00 | 1.80 | 40.00 | 6.80 | single | 6.41% | 10.52 |
| C03F.T005 | C | SNPRC #25567 | 8.38% | 1,000.00 | 13.40 | 40.00 | 29.48 | single | 170.96% | 6.38 |
| C04D.T002 | C | SNPRC #14068 | 0.01% | 8,000.00 | 1.00 | 40.00 | 5.24 | serial | 0.00% | 24.46 |
| C05F.T009 | C | SNPRC #27958 | 17.40% | 800.00 | 22.28 | 40.00 | 36.00 | single | 152.22% | 13.29 |
| C06F.T010 | C | SNPRC #28064 | 4.40% | 1,000.00 | 7.04 | 40.00 | 15.12 | single | 138.62% | 19.02 |
| C07F.C050 | C | Chiou-14-050 | 2.99% | 1,000.00 | 4.78 | 40.00 | 5.24 | single | 14.05% | 7.92 |
| C08D.C050 | C | Chiou-14-050 | 0.82% | 600.00 | 1.00 | 40.00 | 1.02 | serial | 0.00% | 11.08 |
| C09F.B051 | C | BZ06-051 | 3.33% | 500.00 | 2.66 | 40.00 | 2.92 | single | 49.45% | 2.64 |
| C10F.T005 | C | SNPRC #25567 | 12.83% | 1,000.00 | 1.50 | 40.00 | 14.08 | single | 221.02% | 10.31 |
| C11D.C069 | C | Chiou-14-069 | 0.00% | 1,000.00 | 1.00 | 40.00 | 0.76 | serial | 24.47% | 6.81 |
| C12D.B051 | C | BZ06-051 | 0.36% | 600.00 | 1.00 | 40.00 | 1.18 | serial | 14.76% | 2.10 |
| D01D.B220 | D | BZ06-220 | 0.03% | 1,000.00 | 1.00 | 40.00 | 0.70 | serial | - | - |
| D02D.J001 | D | BZ07-001 | 0.02% | 1,000.00 | 1.00 | 40.00 | 2.39 | serial | - | - |
| D03D.J007 | D | BZ07-007 | 0.12% | 947.20 | 1.00 | 40.00 | 0.95 | serial | - | - |
| D04D.J029 | D | BZ07-029 | 0.00% | 1,000.00 | 1.00 | 40.00 | 2.25 | serial | - | - |
| D05D.J032 | D | BZ07-032 | 0.88% | 1,000.00 | 1.00 | 40.00 | 0.97 | serial | - | - |
| D06D.J034 | D | BZ07-034 | 0.47% | 1,000.00 | 1.00 | 40.00 | 2.23 | serial | - | - |
| D07D.J039 | D | BZ07-039 | 0.06% | 1,000.00 | 1.00 | 40.00 | 1.05 | serial | - | - |
| D08D.C057 | D | Chiou-14-057 | 0.24% | 1,000.00 | 1.00 | 40.00 | 0.90 | serial | - | - |
| D09D.C003 | D | Chiou-14-003 | 0.03% | 1,000.00 | 1.00 | 40.00 | 0.88 | serial | - | - |
| D10D.C044 | D | Chiou-14-044 | 0.03% | 913.96 | 1.00 | 40.00 | 0.93 | serial | - | - |
| D11D.C041 | D | Chiou-14-041 | 0.03% | 1,000.00 | 1.00 | 40.00 | 1.31 | serial | - | - |
| D12D.C042 | D | Chiou-14-042 | 0.53% | 1,000.00 | 1.00 | 40.00 | 0.78 | serial | - | - |
| D13D.B221 | D | BZ06-221 | 0.09% | 548.96 | 1.00 | 40.00 | 0.66 | serial | - | - |
| D14D.B227 | D | BZ06-227 | 0.03% | 1,000.00 | 1.00 | 40.00 | 0.52 | serial | - | - |
| D15D.J004 | D | BZ07-004 | 0.39% | 1,000.00 | 1.00 | 40.00 | 0.69 | serial | - | - |
| D16D.J005 | D | BZ07-005 | 0.21% | 1,000.00 | 1.00 | 40.00 | 0.46 | serial | - | - |
| D17D.J030 | D | BZ07-030 | 0.27% | 988.80 | 1.00 | 40.00 | 0.68 | serial | - | - |
| D18D.J035 | D | BZ07-035 | 0.44% | 1,000.00 | 1.00 | 40.00 | 0.70 | serial | - | - |
| D19D.J041 | D | BZ07-041 | 0.44% | 1,000.00 | 1.00 | 40.00 | 0.53 | serial | - | - |
| D20D.C001 | D | Chiou-14-001 | 0.01% | 1,000.00 | 1.00 | 40.00 | 0.64 | serial | - | - |
| D21D.C004 | D | Chiou-14-004 | 0.01% | 1,000.00 | 1.00 | 40.00 | 0.53 | serial | - | - |
| D22D.C065 | D | Chiou-14-065 | 0.08% | 1,000.00 | 1.00 | 40.00 | 0.48 | serial | - | - |

Supplemental Table S2 – continued from previous page
| ID | Lib | Ind | PHB | TD | BV | TV | TY | NE | PHA | NDB |
| --- | --- | --- | --- | --- | --- | --- | --- | --- | --- | --- |
| D23D.C030 | D | Chiou-14-030 | 0.30% | 629.26 | 1.00 | 40.00 | 0.63 | serial | - | - |
| D24D.C039 | D | Chiou-14-039 | 0.29% | 1,000.00 | 1.00 | 40.00 | 0.57 | serial | - | - |
| D25D.B218 | D | BZ06-218 | 0.06% | 893.52 | 1.00 | 40.00 | < 0.40 | serial | - | - |
| D26D.B224 | D | BZ06-224 | 0.03% | 795.70 | 1.00 | 40.00 | < 0.40 | serial | - | - |
| D27D.B225 | D | BZ06-225 | 0.06% | 1,000.00 | 1.00 | 40.00 | < 0.40 | serial | - | - |
| D28D.J042 | D | BZ07-042 | 0.11% | 1,000.00 | 1.00 | 40.00 | < 0.40 | serial | - | - |
| D29D.J045 | D | BZ07-045 | 0.03% | 985.60 | 1.00 | 40.00 | < 0.40 | serial | - | - |
| D30D.J047 | D | BZ07-047 | 0.02% | 1,000.00 | 1.00 | 40.00 | < 0.40 | serial | - | - |
| D31D.J100 | D | BZ07-100 | 0.08% | 1,000.00 | 1.00 | 40.00 | 0.42 | serial | - | - |
| D32D.C036 | D | Chiou-14-036 | 0.37% | 154.76 | 1.00 | 40.00 | 0.42 | serial | - | - |
| D33D.C054 | D | Chiou-14-054 | 0.44% | 1,000.00 | 1.00 | 40.00 | < 0.40 | serial | - | - |
| D34D.C056 | D | Chiou-14-056 | 0.11% | 1,000.00 | 1.00 | 40.00 | < 0.40 | serial | - | - |
| D35D.C058 | D | Chiou-14-058 | 0.57% | 627.80 | 1.00 | 40.00 | < 0.40 | serial | - | - |
| D36D.C059 | D | Chiou-14-059 | 0.90% | 383.98 | 1.00 | 40.00 | < 0.40 | serial | - | - |
| D37D.H003 | D | Chiou-15-003 | 0.10% | 220.46 | 1.00 | 40.00 | < 0.40 | serial | - | - |
| D38D.H004 | D | Chiou-15-004 | 3.11% | 186.88 | 1.00 | 40.00 | < 0.40 | serial | - | - |
| D39D.H005 | D | Chiou-15-005 | 0.11% | 511.00 | 1.00 | 40.00 | < 0.40 | serial | - | - |
| D40D.C005 | D | Chiou-14-005 | 2.41% | 182.50 | 1.00 | 40.00 | < 0.40 | serial | - | - |

**Supplemental Table S3:** Library preparation and sequence mapping results. Key: ID, experiment ID (see Supplemental Table S2); T, tissue type; PHB, percent host DNA before; TD, total DNA used (ng); PID, pool ID; PC, total number of PCR amplification cycles; TR, total number of sequencing reads; RM, number of reads mapping to the baboon reference genome (papAnu2); PRM, percentage of reads mapping to the baboon reference genome (papAnu2).

| ID | T | PHB | TD | PID | PC | TR | RM | PRM |
| --- | --- | --- | --- | --- | --- | --- | --- | --- |
| A01F.T002 | feces | 0.1500% | 83.00 | A1 | 24 | 264,158 | 9,856 | 3.73% |
| A02F.T005 | feces | 4.4000% | 200.00 | A1 | 24 | 2,607,006 | 1,892,463 | 72.59% |
| A03F.T007 | feces | 0.0000% | 191.00 | A1 | 24 | 2,040,002 | 15,224 | 0.75% |
| A04F.T009 | feces | 13.5900% | 200.00 | A2 | 24 | 4,104,470 | 3,334,314 | 81.24% |
| A05F.B051 | feces | 1.5900% | 122.00 | A2 | 24 | 916,680 | 257,714 | 28.11% |
| A06F.B053 | feces | 1.1400% | 120.00 | A2 | 24 | 591,626 | 146,285 | 24.73% |
| A07B.T002 | blood | - | 200.00 | A3 | 24 | 2,683,338 | 2,384,501 | 88.86% |
| A08B.T005 | blood | - | 200.00 | A3 | 24 | 745,504 | 681,730 | 91.45% |
| A09B.T007 | blood | - | 200.00 | A4 | 24 | 2,128,974 | 1,886,608 | 88.62% |
| A10B.T009 | blood | - | 200.00 | A4 | 24 | 1,189,390 | 1,087,777 | 91.46% |
| B01F.T001 | feces | 3.5500% | 60.06 | B1 | 24 | 4,882,156 | 1,652,610 | 33.85% |
| B02F.T002 | feces | 0.1500% | 13.27 | B1 | 24 | 2,507,424 | 111,206 | 4.44% |
| B03F.T007 | feces | 0.0000% | 22.11 | B1 | 24 | 2,461,390 | 27,185 | 1.10% |
| B04F.T010 | feces | 9.8700% | 65.01 | B1 | 24 | 5,444,582 | 3,184,008 | 58.48% |
| B05F.C030 | feces | 1.2400% | 33.39 | B1 | 24 | 619,862 | 34,761 | 5.61% |
| B06F.C050 | feces | 2.4000% | 11.62 | B1 | 24 | 2,123,294 | 584,719 | 27.54% |
| B07F.C065 | feces | 0.7600% | 20.09 | B1 | 24 | 1,241,912 | 44,732 | 3.60% |
| B08F.C069 | feces | 0.4500% | 13.93 | B1 | 24 | 1,203,334 | 23,642 | 1.96% |
| B09F.B066 | feces | 4.8500% | 34.30 | B1 | 24 | 1,563,428 | 581,005 | 37.16% |
| B10F.B148 | feces | 0.6800% | 8.68 | B1 | 24 | 501,188 | 62,126 | 12.40% |
| B11B.T001 | blood | - | 196.50 | B2 | 24 | 2,216,126 | 1,819,922 | 82.12% |
| B12B.T010 | blood | - | 201.50 | B2 | 24 | 2,092,674 | 1,780,702 | 85.09% |
| C01F.T001 | feces | 4.3300% | 17.68 | C1 | 20 | 1,782,002 | 1,188,499 | 66.69% |
| C02F.T002 | feces | 1.1200% | 5.78 | C2 | 24 | 1,290,578 | 333,245 | 25.82% |
| C03F.T005 | feces | 8.3800% | 25.06 | C1 | 20 | 1,871,868 | 1,316,267 | 70.32% |
| C04D.T002 | feces | 0.0100% | 3.41 | C3 | 26 | 1,841,762 | 163,819 | 8.89% |
| C05F.T009 | feces | 17.4000% | 30.60 | C1 | 20 | 3,116,288 | 2,345,065 | 75.25% |
| C06F.T010 | feces | 4.4000% | 12.85 | C1 | 20 | 1,469,246 | 967,950 | 65.88% |
| C07F.C050 | feces | 2.9900% | 4.45 | C2 | 24 | 3,570,776 | 1,845,128 | 51.67% |
| C08D.C050 | feces | 0.8200% | 0.67 | C4 | 26 | 2,059,728 | 737,230 | 35.79% |
| C09F.B051 | feces | 3.3300% | 2.48 | C2 | 24 | 1,816,378 | 918,754 | 50.58% |
| C10F.T005 | feces | 12.8300% | 11.97 | C1 | 20 | 2,068,478 | 1,475,254 | 71.32% |
| C11D.C069 | feces | 0.0049% | 0.49 | C5 | 26 | 1,514,706 | 195,365 | 12.90% |
| C12D.B051 | feces | 0.3600% | 0.77 | C6 | 26 | 1,896,580 | 1,035,140 | 54.58% |
| D01D.B220 | feces | 0.0289% | 0.53 | D1 | 22 | 5,875,038 | 558,679 | 9.51% |
| D02D.J001 | feces | 0.0158% | 1.79 | D1 | 22 | 1,449,446 | 47,799 | 3.30% |
| D03D.J007 | feces | 0.1220% | 0.71 | D1 | 22 | 3,243,182 | 760,274 | 23.44% |
| D04D.J029 | feces | 0.0030% | 1.69 | D1 | 22 | 1,542,546 | 22,534 | 1.46% |
| D05D.J032 | feces | 0.8793% | 0.73 | D1 | 22 | 9,398,314 | 6,856,889 | 72.96% |
| D06D.J034 | feces | 0.4713% | 1.67 | D1 | 22 | 2,656,920 | 1,313,120 | 49.42% |
| D07D.J039 | feces | 0.0629% | 0.79 | D1 | 22 | 2,230,514 | 609,824 | 27.34% |
| D08D.C057 | feces | 0.2374% | 0.67 | D1 | 22 | 24,351,758 | 9,032,717 | 37.09% |
| D09D.C003 | feces | 0.0339% | 0.66 | D1 | 22 | 4,453,940 | 140,853 | 3.16% |
| D10D.C044 | feces | 0.0292% | 0.70 | D1 | 22 | 2,137,704 | 124,844 | 5.84% |
| D11D.C041 | feces | 0.0271% | 0.98 | D1 | 22 | 1,140,122 | 96,922 | 8.50% |
| D12D.C042 | feces | 0.5310% | 0.59 | D1 | 22 | 10,202,338 | 4,952,523 | 48.54% |
| D13D.B221 | feces | 0.0894% | 0.49 | D2 | 22 | 5,147,652 | 982,161 | 19.08% |
| D14D.B227 | feces | 0.0309% | 0.39 | D2 | 22 | 1,828,496 | 188,056 | 10.28% |
| D15D.J004 | feces | 0.3858% | 0.52 | D2 | 22 | 3,606,424 | 1,619,089 | 44.89% |

Supplemental Table S3 – continued from previous page
| ID | T | PHB | TD | PID | PC | TR | RM | PRM |
| --- | --- | --- | --- | --- | --- | --- | --- | --- |
| D16D.J005 | feces | 0.2120% | 0.35 | D2 | 22 | 4,681,458 | 1,595,305 | 34.08% |
| D17D.J030 | feces | 0.2748% | 0.51 | D2 | 22 | 9,260,376 | 4,298,954 | 46.42% |
| D18D.J035 | feces | 0.4370% | 0.52 | D2 | 22 | 14,641,030 | 7,429,804 | 50.75% |
| D19D.J041 | feces | 0.4441% | 0.40 | D2 | 22 | 2,734,646 | 1,830,636 | 66.94% |
| D20D.C001 | feces | 0.0079% | 0.48 | D2 | 22 | 3,011,748 | 95,667 | 3.18% |
| D21D.C004 | feces | 0.0088% | 0.40 | D2 | 22 | 2,685,536 | 97,831 | 3.64% |
| D22D.C065 | feces | 0.0770% | 0.36 | D2 | 22 | 6,516,948 | 954,911 | 14.65% |
| D23D.C030 | feces | 0.2962% | 0.47 | D2 | 22 | 4,200,672 | 1,854,549 | 44.15% |
| D24D.C039 | feces | 0.2900% | 0.43 | D2 | 22 | 15,175,272 | 5,263,567 | 34.69% |
| D25D.B218 | feces | 0.0581% | 0.23 | D3 | 26 | 2,624,536 | 311,121 | 11.85% |
| D26D.B224 | feces | 0.0307% | 0.23 | D3 | 26 | 3,581,376 | 246,941 | 6.90% |
| D27D.B225 | feces | 0.0573% | 0.23 | D3 | 26 | 16,512,734 | 1,228,293 | 7.44% |
| D28D.J042 | feces | 0.1062% | 0.23 | D3 | 26 | 6,967,098 | 1,135,899 | 16.30% |
| D29D.J045 | feces | 0.0262% | 0.23 | D3 | 26 | 2,307,634 | 124,309 | 5.39% |
| D30D.J047 | feces | 0.0213% | 0.23 | D3 | 26 | 5,640,310 | 519,703 | 9.21% |
| D31D.J100 | feces | 0.0793% | 0.31 | D3 | 26 | 1,219,418 | 83,022 | 6.81% |
| D32D.C036 | feces | 0.3708% | 0.32 | D3 | 26 | 7,119,672 | 4,402,290 | 61.83% |
| D33D.C054 | feces | 0.4415% | 0.23 | D4 | 22 | 7,013,280 | 4,484,858 | 63.95% |
| D34D.C056 | feces | 0.1098% | 0.23 | D4 | 22 | 7,949,918 | 2,748,447 | 34.57% |
| D35D.C058 | feces | 0.5733% | 0.23 | D4 | 22 | 10,539,604 | 4,223,730 | 40.07% |
| D36D.C059 | feces | 0.8954% | 0.23 | D4 | 22 | 5,962,728 | 3,090,413 | 51.83% |
| D37D.H003 | feces | 0.1017% | 0.23 | D4 | 22 | 1,418,168 | 74,131 | 5.23% |
| D38D.H004 | feces | 3.1094% | 0.23 | D4 | 22 | 9,161,532 | 6,739,523 | 73.56% |
| D39D.H005 | feces | 0.1113% | 0.23 | D4 | 22 | 1,784,298 | 585,250 | 32.80% |
| D40D.C005 | feces | 2.4120% | 0.23 | D4 | 22 | 2,621,020 | 1,611,725 | 61.49% |
| E01B.T001 | blood | - | 200.00 | E1 | 12 | 2,326,792 | 2,027,296 | 87.13% |
| E02B.T002 | blood | - | 200.00 | E1 | 12 | 1,249,950 | 1,095,499 | 87.64% |
| E03B.T005 | blood | - | 200.00 | E2 | 12 | 4,812,938 | 4,192,408 | 87.11% |
| E04B.T009 | blood | - | 200.00 | E1 | 12 | 1,986,292 | 1,746,868 | 87.95% |
| E05B.T010 | blood | - | 200.00 | E2 | 12 | 4,091,500 | 3,597,699 | 87.93% |

**Supplemental Table S4:** DNA samples used for controlled experiments. Artificial “fecal” DNA was prepared by manually mixing DNA samples in controlled proportions. Artificial methylated DNA was also prepared using amplicons of lambda phage DNA (with known sequence) and methyltransferase enzymes with specific recognition sites. 5,012 bp amplicons were prepared using the primers `/5Biosg/GTTCTGCACTGACAGATTAAAACTCG` and CTGCTCATTAATATACTTCTGGGTTCC, 15,089 bp amplicons were prepared using the primers `/5Biosg/GAGTGAATATATCGAACAGTCAGG` and GTGTCATATTTCACTTCCGTACC, and 10,144 bp amplicons were prepared using the primers `/5BiosG/ATAAAGATGAGACGCTGGAGTACA` and GCGATAACCAGGTAAAATTTTCCG. Key: ID, prepared DNA sample ID; DNA1, input DNA sample 1; DNA2, input DNA sample 2; PH, percentage of “host” (baboon) DNA; PB, percentage of bacterial DNA; L, length of DNA amplicon; Enz, methyltransferase enzyme(s) used; MD, CpG methylation density.

| ID | DNA1 | DNA2 | PH | PB | L | Enz | MD |
| --- | --- | --- | --- | --- | --- | --- | --- |
| PB01 | K12 *E. coli* | none | 0.0% | 100.00% | - | - | - |
| PB02 | ATCC 11303 *E. coli* | none | 0.0% | 100.00% | - | - | - |
| PH01 | Baboon blood | none | 100.0% | 0.0% | - | - | - |
| PH02 | Baboon liver | none | 100.0% | 0.0% | - | - | - |
| AF01 | Baboon blood | K12 *E. coli* | 2.0% | 98.0% | - | - | - |
| AF02 | Baboon blood | K12 *E. coli* | 0.2% | 99.8% | - | - | - |
| AF03 | Baboon blood | K12 *E. coli* | 50.0% | 50.0% | - | - | - |
| AF04 | Baboon blood | K12 *E. coli* | 2.0% | 98.0% | - | - | - |
| AF05 | Baboon blood | K12 *E. coli* | 50.0% | 50.0% | - | - | - |
| AF06 | Baboon blood | K12 *E. coli* | 5.0% | 95.0% | - | - | - |
| AF07 | Baboon blood | K12 *E. coli* | 10.0% | 90.0% | - | - | - |
| AF08 | Baboon blood | ATCC 11303 *E. coli* | 2.0% | 98.0% | - | - | - |
| AF09 | Baboon liver | ATCC 11303 *E. coli* | 2.0% | 98.0% | - | - | - |
| AF10 | Baboon liver | ATCC 11303 *E. coli* | 50.0% | 50.0% | - | - | - |
| AF11 | Baboon liver | ATCC 11303 *E. coli* | 0.5% | 99.5% | - | - | - |
| AF12 | Baboon liver | ATCC 11303 *E. coli* | 2.0% | 98.0% | - | - | - |
| AF13 | Baboon liver | ATCC 11303 *E. coli* | 5.0% | 95.0% | - | - | - |
| AF14 | Baboon liver | ATCC 11303 *E. coli* | 2.0% | 98.0% | - | - | - |
| AF15 | Baboon liver | ATCC 11303 *E. coli* | 0.5% | 99.5% | - | - | - |
| AF16 | Baboon liver | ATCC 11303 *E. coli* | 2.0% | 98.0% | - | - | - |
| AF17 | Baboon liver | ATCC 11303 *E. coli* | 5.0% | 95.0% | - | - | - |
| AF18 | Baboon liver | ATCC 11303 *E. coli* | 0.5% | 99.5% | - | - | - |
| CD01 | Lambda cl857 phage | none | 0.0% | 0.0% | 5,012 | *HhaI* | 3.6 |
| CD02 | Lambda cl857 phage | none | 0.0% | 0.0% | 5,012 | - | 0.0 |
| CD03 | Lambda cl857 phage | none | 0.0% | 0.0% | 5,012 | *HhaI* | 3.6 |
| CD04 | Lambda cl857 phage | none | 0.0% | 0.0% | 5,012 | *HhaI* + *HpaII* | 7.2 |
| CD05 | Lambda cl857 phage | none | 0.0% | 0.0% | 5,012 | *HhaI* | 3.6 |
| CD06 | Lambda cl857 phage | none | 0.0% | 0.0% | 15,089 | *HhaI* | 6.9 |
| CD07 | Lambda cl857 phage | none | 0.0% | 0.0% | 10,144 | *HhaI* | 6.3 |
| CD08 | Lambda cl857 phage | none | 0.0% | 0.0% | 10,144 | *HhaI* + *HpaII* | 17.7 |

**Supplemental Table S5:** Controlled DNA enrichment experiments. DNA enrichment was simulated from artificial “fecal” samples. In some cases, additional DNA was included. A number of variables described in Supplemental Protocol were tuned to evaluate their impact on enrichment results (see Supplemental Table S6). Key: ID, experiment ID; SID, experiment set ID; DNA1, input DNA sample 1 (see Supplemental Table S4 or this table); TD1, total amount of sample 1 (ng); PH, percentage of “host” (baboon) DNA in sample 1; DNA2, input DNA sample 2 (see Supplemental Table S4); TD2, total amount of sample 2 (ng); BV, volume of protein A beads used (µl); PV, volume of MBD-Fc protein used (µl); NC, NaCl concentration of reaction (µM); TV, total volume of reaction (µl); NW, number of washes; NCW, NaCl concentration of each wash (µM); WV, volume of each wash (µl); EM, Elution method. For elutions in TE, proteinase K was added at a ratio of 1 µl proteinase K to 10 µl 1X TE.

| ID | SID | DNA1 | TD1 | PH | DNA2 | TD2 | BV | PV | NC | TV | NW | NCW | WV | EM |
| --- | --- | --- | --- | --- | --- | --- | --- | --- | --- | --- | --- | --- | --- | --- |
| X001 | S01 | AF01 | 1,000.00 | 2.0% | - | 0.00 | 160.0 | 16.00 | 150 | 166.20 | 0 | - | - | 150 μl TE |
| X002 | S01 | AF01 | 2,000.00 | 2.0% | - | 0.00 | 160.0 | 16.00 | 150 | 172.40 | 0 | - | - | 150 μl TE |
| X003 | S01 | AF02 | 1,000.00 | 0.2% | - | 0.00 | 160.0 | 16.00 | 150 | 164.00 | 0 | - | - | 150 μl TE |
| X004 | S01 | AF01 | 1,000.00 | 2.0% | - | 0.00 | 320.0 | 32.00 | 150 | 326.20 | 0 | - | - | 150 μl TE |
| X005 | S01 | AF01 | 1,000.00 | 2.0% | - | 0.00 | 80.0 | 8.00 | 150 | 86.20 | 0 | - | - | 150 μl TE |
| X006 | S01 | AF01 | 1,000.00 | 2.0% | - | 0.00 | 40.0 | 4.00 | 150 | 46.20 | 0 | - | - | 150 μl TE |
| X007 | S02 | PB01 | 1,000.00 | 0.0% | - | 0.00 | 160.0 | 16.00 | 150 | 163.60 | 0 | - | - | 150 μl TE |
| X008 | S02 | AF01 | 1,000.00 | 2.0% | - | 0.00 | 16.0 | 1.60 | 150 | 22.20 | 0 | - | - | 150 μl TE |
| X009 | S02 | AF03 | 40.00 | 50.0% | - | 0.00 | 80.0 | 8.00 | 150 | 130.00 | 0 | - | - | 150 μl TE |
| X010 | S02 | AF03 | 40.00 | 50.0% | - | 0.00 | 40.0 | 4.00 | 150 | 90.00 | 0 | - | - | 150 μl TE |
| X011 | S02 | AF03 | 40.00 | 50.0% | - | 0.00 | 16.0 | 1.60 | 150 | 66.00 | 0 | - | - | 150 μl TE |
| X012 | S02 | AF03 | 40.00 | 50.0% | - | 0.00 | 8.0 | 0.80 | 150 | 58.00 | 0 | - | - | 150 μl TE |
| X013 | S03 | PB01 | 1,000.00 | 0.0% | - | 0.00 | 160.0 | 0.00 | 150 | 163.60 | 0 | - | - | 150 μl TE |
| X014 | S03 | PB01 | 1,000.00 | 0.0% | - | 0.00 | 40.0 | 0.00 | 150 | 47.20 | 0 | - | - | 150 μl TE |
| X015 | S03 | PB01 | 200.00 | 0.0% | - | 0.00 | 40.0 | 0.00 | 150 | 47.20 | 0 | - | - | 150 μl TE |
| X016 | S03 | PB01 | 1,000.00 | 0.0% | - | 0.00 | 40.0 | 32.00 | 150 | 47.20 | 0 | - | - | 150 μl TE |
| X017 | S04 | AF04 | 1,000.00 | 2.0% | - | 0.00 | 1.0 | 0.10 | 150 | 7.20 | 0 | - | - | 150 μl TE |
| X018 | S04 | PB01 | 1,000.00 | 0.0% | - | 0.00 | 40.0 | 4.00 | 150 | 43.60 | 0 | - | - | 150 μl TE |
| X019 | S04 | PB01 | 1,000.00 | 0.0% | - | 0.00 | 40.0 | 4.00 | 300 | 58.20 | 0 | - | - | 150 μl TE |
| X020 | S04 | AF05 | 1,000.00 | 50.0% | - | 0.00 | 40.0 | 4.00 | 150 | 105.80 | 0 | - | - | 150 μl TE |
| X021 | S05 | AF04 | 1,000.00 | 2.0% | - | 0.00 | 40.0 | 4.00 | 150 | 46.20 | 0 | - | - | 150 μl TE |
| X022 | S05 | AF04 | 1,000.00 | 2.0% | - | 0.00 | 40.0 | 4.00 | 150 | 46.20 | 0 | - | - | 150 μl TE |
| X023 | S05 | X001 | 29.16 | 51.0% | - | 0.00 | 40.0 | 4.00 | 150 | 78.60 | 0 | - | - | 150 μl TE |
| X024 | S05 | X006 | 21.48 | 44.1% | - | 0.00 | 40.0 | 4.00 | 150 | 78.40 | 0 | - | - | 150 μl TE |
| X025 | S06 | AF04 | 1,000.00 | 2.0% | CD01 | 500.00 | 40.0 | 4.00 | 150 | 48.90 | 0 | - | - | 150 μl TE |
| X026 | S06 | AF06 | 1,000.00 | 5.0% | - | 0.00 | 40.0 | 4.00 | 150 | 49.90 | 0 | - | - | 150 μl TE |
| X027 | S06 | AF07 | 1,000.00 | 10.0% | - | 0.00 | 40.0 | 4.00 | 150 | 56.10 | 0 | - | - | 150 μl TE |
| X028 | S06 | AF04 | 1,000.00 | 2.0% | CD02 | 500.00 | 40.0 | 4.00 | 150 | 48.70 | 0 | - | - | 150 μl TE |
| X029 | S07 | AF04 | 1,000.00 | 2.0% | CD01 | 2,500.00 | 40.0 | 4.00 | 150 | 58.70 | 0 | - | - | 150 μl TE |
| X030 | S07 | AF04 | 1,000.00 | 2.0% | - | 0.00 | 1.0 | 0.10 | 150 | 7.20 | 0 | - | - | 150 μl TE |
| X031 | S07 | AF04 | 1,000.00 | 2.0% | - | 0.00 | 1.0 | 0.10 | 150 | 100.00 | 0 | - | - | 150 μl TE |
| X032 | S07 | AF04 | 1,000.00 | 2.0% | CD01 | 500.00 | 1.0 | 0.10 | 150 | 9.70 | 0 | - | - | 150 μl TE |
| X033 | S08 | AF08 | 1,000.00 | 2.0% | - | 0.00 | 8.0 | 0.80 | 150 | 13.00 | 0 | - | - | 150 μl TE |
| X034 | S08 | AF08 | 1,000.00 | 2.0% | - | 0.00 | 4.0 | 0.40 | 150 | 9.00 | 0 | - | - | 150 μl TE |
| X035 | S08 | AF08 | 1,000.00 | 2.0% | - | 0.00 | 2.0 | 0.20 | 150 | 7.00 | 0 | - | - | 150 μl TE |
| X036 | S08 | AF08 | 1,000.00 | 2.0% | - | 0.00 | 0.5 | 0.05 | 150 | 6.00 | 0 | - | - | 150 μl TE |
| X037 | S08 | PB02 | 980.00 | 0.0% | - | 0.00 | 1.0 | 0.10 | 150 | 7.20 | 0 | - | - | 150 μl TE |
| X038 | S08 | PH01 | 20.00 | 100.0% | - | 0.00 | 1.0 | 0.10 | 150 | 7.20 | 0 | - | - | 150 μl TE |
| X039 | S09 | AF08 | 1,000.00 | 2.0% | CD03 | 1,000.00 | 40.0 | 4.00 | 150 | 56.30 | 0 | - | - | 150 μl TE |
| X040 | S09 | AF08 | 1,000.00 | 2.0% | CD03 | 500.00 | 40.0 | 4.00 | 150 | 50.60 | 0 | - | - | 150 μl TE |
| X041 | S10 | AF09 | 1,000.00 | 2.0% | CD04 | 500.00 | 40.0 | 4.00 | 150 | 80.00 | 0 | - | - | 150 μl TE |
| X042 | S10 | AF09 | 1,000.00 | 2.0% | CD04 | 500.00 | 16.0 | 1.60 | 150 | 32.00 | 0 | - | - | 150 μl TE |

Supplemental Table S5 – continued from previous page
| ID | EID | DNA1 | TD1 | PH | DNA2 | TD2 | BV | PV | NC | TV | NW | NCW | WV | EM |
| --- | --- | --- | --- | --- | --- | --- | --- | --- | --- | --- | --- | --- | --- | --- |
| X043 | S11 | PH02 | 1,000.00 | 100.0% | - | 0.00 | 1.0 | 0.10 | 150 | 12.13 | 0 | - | - | 150 µl TE |
| X044 | S11 | PH02 | 2,000.00 | 100.0% | - | 0.00 | 1.0 | 0.10 | 150 | 23.36 | 0 | - | - | 150 µl TE |
| X045 | S11 | PH02 | 1,000.00 | 100.0% | - | 0.00 | 2.0 | 0.20 | 150 | 13.13 | 0 | - | - | 150 µl TE |
| X046 | S11 | PH02 | 1,000.00 | 100.0% | - | 0.00 | 4.0 | 0.40 | 150 | 15.13 | 0 | - | - | 150 µl TE |
| X047 | S11 | PH02 | 1,000.00 | 100.0% | - | 0.00 | 8.0 | 0.80 | 150 | 19.13 | 0 | - | - | 150 µl TE |
| X048 | S11 | PH02 | 1,000.00 | 100.0% | - | 0.00 | 16.0 | 1.60 | 150 | 27.13 | 0 | - | - | 150 µl TE |
| X049 | S12 | PH02 | 112.00 | 100.0% | - | 0.00 | 1.0 | 0.10 | 150 | 2.25 | 0 | - | - | 150 µl TE |
| X050 | S12 | PH02 | 112.00 | 100.0% | - | 0.00 | 1.0 | 0.10 | 150 | 2.25 | 0 | - | - | 40 µl TE |
| X051 | S12 | PH02 | 112.00 | 100.0% | - | 0.00 | 1.0 | 0.10 | 150 | 2.25 | 0 | - | - | 150 µl TE |
| X052 | S12 | PH02 | 112.00 | 100.0% | - | 0.00 | 1.0 | 0.10 | 150 | 2.25 | 0 | - | - | 60 µl TE |
| X053 | S13 | PB02 | 1,000.00 | 0.0% | CD05 | 1,000.00 | 40.0 | 4.00 | 150 | 80.00 | 0 | - | - | 150 µl TE |
| X054 | S13 | AF10 | 2,000.00 | 50.0% | - | 0.00 | 40.0 | 4.00 | 150 | 80.00 | 0 | - | - | 150 µl TE |
| X055 | S14 | PB02 | 1,000.00 | 0.0% | CD06 | 1,000.00 | 40.0 | 4.00 | 150 | 80.00 | 0 | - | - | 150 µl TE |
| X056 | S14 | AF11 | 1,000.00 | 0.5% | CD06 | 1,000.00 | 40.0 | 4.00 | 150 | 80.00 | 0 | - | - | 150 µl TE |
| X057 | S14 | AF12 | 1,000.00 | 2.0% | CD06 | 1,000.00 | 40.0 | 4.00 | 150 | 80.00 | 0 | - | - | 150 µl TE |
| X058 | S14 | AF13 | 1,000.00 | 5.0% | CD06 | 1,000.00 | 40.0 | 4.00 | 150 | 80.00 | 0 | - | - | 150 µl TE |
| X059 | S15 | PB02 | 1,000.00 | 0.0% | - | 0.00 | 40.0 | 4.00 | 150 | 80.00 | 1 | 150 | 80 | see Supplemental Table S7 |
| X060 | S15 | PH02 | 250.00 | 100.0% | - | 0.00 | 40.0 | 4.00 | 150 | 80.00 | 1 | 150 | 80 | see Supplemental Table S7 |
| X061 | S16 | AF14 | 1,000.00 | 2.0% | - | 0.00 | 40.0 | 4.00 | 150 | 80.00 | 0 | - | - | 2 M NaCl |
| X062 | S16 | AF14 | 1,000.00 | 2.0% | - | 0.00 | 40.0 | 1.00 | 150 | 80.00 | 0 | - | - | 2 M NaCl |
| X063 | S16 | AF14 | 1,000.00 | 2.0% | CD07 | 1,000.00 | 40.0 | 4.00 | 150 | 80.00 | 0 | - | - | 2 M NaCl |
| X064 | S16 | AF14 | 1,000.00 | 2.0% | CD07 | 1,000.00 | 40.0 | 4.00 | 150 | 80.00 | 0 | - | - | 2 M NaCl |
| X065 | S16 | AF14 | 1,000.00 | 2.0% | - | 0.00 | 40.0 | 4.00 | 200 | 80.00 | 0 | - | - | 2 M NaCl |
| X066 | S16 | AF14 | 1,000.00 | 2.0% | - | 0.00 | 40.0 | 4.00 | 150 | 80.00 | 1 | 200 | 200 | 2 M NaCl |
| X067 | S17 | AF14 | 1,000.00 | 2.0% | - | 0.00 | 40.0 | 4.00 | 150 | 80.00 | 0 | - | - | 150 µl TE |
| X068 | S17 | AF14 | 1,000.00 | 2.0% | - | 0.00 | 40.0 | 4.00 | 150 | 80.00 | 0 | - | - | 2 M NaCl |
| X069 | S17 | AF14 | 1,000.00 | 2.0% | - | 0.00 | 40.0 | 0.25 | 150 | 80.00 | 0 | - | - | 2 M NaCl |
| X070 | S17 | AF14 | 1,000.00 | 2.0% | - | 0.00 | 40.0 | 0.25 | 150 | 80.00 | 1 | 150 | 100 | 2 M NaCl |
| X071 | S17 | AF14 | 1,000.00 | 2.0% | - | 0.00 | 40.0 | 0.25 | 150 | 80.00 | 1 | 200 | 100 | 2 M NaCl |
| X072 | S17 | AF14 | 1,000.00 | 2.0% | CD07 | 1,000.00 | 40.0 | 0.25 | 150 | 80.00 | 1 | 200 | 100 | 2 M NaCl |
| X073 | S18 | AF14 | 1,000.00 | 2.0% | - | 0.00 | 40.0 | 1.00 | 150 | 80.00 | 1 | 150 | 100 | 2 M NaCl |
| X074 | S18 | AF14 | 1,000.00 | 2.0% | - | 0.00 | 40.0 | 1.00 | 150 | 80.00 | 1 | 200 | 100 | 2 M NaCl |
| X075 | S18 | AF14 | 2,000.00 | 2.0% | - | 0.00 | 40.0 | 1.00 | 150 | 80.00 | 1 | 150 | 100 | 2 M NaCl |
| X076 | S19 | AF15 | 1,000.00 | 0.5% | - | 0.00 | 40.0 | 1.00 | 150 | 80.00 | 1 | 150 | 100 | 2 M NaCl |
| X077 | S19 | AF15 | 1,000.00 | 0.5% | - | 0.00 | 40.0 | 1.00 | 150 | 80.00 | 2 | 150 | 100 | 2 M NaCl |
| X078 | S19 | AF15 | 1,000.00 | 0.5% | - | 0.00 | 40.0 | 1.00 | 150 | 80.00 | 3 | 150 | 100 | 2 M NaCl |
| X079 | S20 | AF15 | 1,000.00 | 0.5% | - | 0.00 | 40.0 | 1.00 | 150 | 80.00 | 1 | 150 | 80 | 2 M NaCl |
| X080 | S20 | AF15 | 1,000.00 | 0.5% | - | 0.00 | 40.0 | 1.00 | 150 | 80.00 | 1 | 150 | 40 | 2 M NaCl |
| X081 | S20 | AF15 | 1,000.00 | 0.5% | CD07 | 40.00 | 40.0 | 1.00 | 150 | 80.00 | 1 | 150 | 80 | 2 M NaCl |
| X082 | S20 | PB02 | 1,000.00 | 0.0% | CD07 | 40.00 | 40.0 | 1.00 | 150 | 80.00 | 1 | 150 | 80 | 2 M NaCl |
| X083 | S21 | AF14 | 1,000.00 | 2.0% | - | 0.00 | 40.0 | 1.00 | 150 | 80.00 | 1 | 150 | 80 | 2 M NaCl |
| X084 | S21 | AF14 | 1,000.00 | 2.0% | - | 0.00 | 40.0 | 1.00 | 150 | 80.00 | 1 | 150 | 200 | 2 M NaCl |
| X085 | S21 | AF14 | 1,000.00 | 2.0% | - | 0.00 | 40.0 | 4.00 | 150 | 80.00 | 1 | 150 | 80 | 2 M NaCl |
| X086 | S21 | AF15 | 1,000.00 | 0.5% | - | 0.00 | 40.0 | 4.00 | 150 | 80.00 | 1 | 150 | 80 | 2 M NaCl |
| X087 | S21 | AF15 | 1,000.00 | 0.5% | CD08 | 20.00 | 40.0 | 4.00 | 150 | 80.00 | 1 | 150 | 80 | 2 M NaCl |
| X088 | S21 | PB02 | 1,000.00 | 0.0% | CD08 | 20.00 | 40.0 | 4.00 | 150 | 80.00 | 1 | 150 | 80 | 2 M NaCl |
| X089 | S22 | AF15 | 1,000.00 | 0.5% | - | 0.00 | 40.0 | 1.00 | 150 | 80.00 | 1 | 150 | 80 | 2 M NaCl |
| X090 | S22 | AF15 | 1,000.00 | 0.5% | CD08 | 20.00 | 40.0 | 1.00 | 150 | 80.00 | 1 | 150 | 80 | 2 M NaCl |
| X091 | S22 | AF14 | 1,000.00 | 2.0% | CD08 | 20.00 | 40.0 | 1.00 | 150 | 80.00 | 1 | 150 | 80 | 2 M NaCl |
| X092 | S22 | PB02 | 1,000.00 | 0.0% | CD08 | 20.00 | 40.0 | 1.00 | 150 | 80.00 | 1 | 150 | 80 | 2 M NaCl |
| X093 | S23 | AF15 | 1,000.00 | 0.5% | - | 0.00 | 40.0 | 0.25 | 150 | 80.00 | 1 | 150 | 100 | 2 M NaCl |
| X094 | S23 | AF15 | 1,000.00 | 0.5% | - | 0.00 | 40.0 | 0.25 | 150 | 80.00 | 2 | 150 | 100 | 2 M NaCl |
| X095 | S23 | AF15 | 1,000.00 | 0.5% | - | 0.00 | 40.0 | 0.25 | 150 | 80.00 | 3 | 150 | 100 | 2 M NaCl |
| X096 | S23 | AF15 | 1,000.00 | 0.5% | - | 0.00 | 40.0 | 0.25 | 150 | 80.00 | 4 | 150 | 100 | 2 M NaCl |
| X097 | S24 | AF15 | 1,000.00 | 0.5% | - | 0.00 | 40.0 | 0.25 | 150 | 80.00 | 1 | 150 | 100 | 2 M NaCl |

Supplemental Table S5 – continued from previous page
| ID | EID | DNA1 | TD1 | PH | DNA2 | TD2 | BV | PV | NC | TV | NW | NCW | WV | EM |
| --- | --- | --- | --- | --- | --- | --- | --- | --- | --- | --- | --- | --- | --- | --- |
| X098 | S24 | AF15 | 1,000.00 | 0.5% | - | 0.00 | 2.5 | 0.25 | 150 | 80.00 | 1 | 150 | 100 | 2 M NaCl |
| X099 | S24 | AF16 | 1,000.00 | 2.0% | - | 0.00 | 40.0 | 0.25 | 150 | 80.00 | 1 | 150 | 100 | 2 M NaCl |
| X100 | S24 | AF17 | 1,000.00 | 5.0% | - | 0.00 | 40.0 | 0.25 | 150 | 80.00 | 1 | 150 | 100 | 2 M NaCl |
| X101 | S25 | AF15 | 1,000.00 | 0.5% | - | 0.00 | 2.5 | 0.25 | 150 | 40.00 | 1 | 150 | 100 | 2 M NaCl |
| X102 | S25 | AF16 | 1,000.00 | 2.0% | - | 0.00 | 10.0 | 1.00 | 150 | 40.00 | 1 | 150 | 100 | 2 M NaCl |
| X103 | S25 | AF17 | 1,000.00 | 5.0% | - | 0.00 | 25.0 | 2.50 | 150 | 40.00 | 1 | 150 | 100 | 2 M NaCl |
| X104 | S25 | AF15 | 1,000.00 | 0.5% | - | 0.00 | 2.5 | 0.25 | 150 | 40.00 | 1 | 150 | 40 | 2 M NaCl |
| X105 | S25 | AF16 | 1,000.00 | 2.0% | - | 0.00 | 3.2 | 0.32 | 150 | 40.00 | 1 | 150 | 100 | 2 M NaCl |
| X106 | S25 | AF15 | 1,000.00 | 0.5% | - | 0.00 | 0.8 | 0.08 | 150 | 40.00 | 1 | 150 | 100 | 2 M NaCl |
| X107 | S26 | AF15 | 1,000.00 | 0.5% | - | 0.00 | 2.5 | 0.25 | 150 | 40.00 | 1 | 150 | 100 | 2 M NaCl |
| X108 | S26 | AF15 | 1,000.00 | 0.5% | - | 0.00 | 0.8 | 0.08 | 150 | 40.00 | 1 | 150 | 100 | 2 M NaCl |
| X109 | S26 | AF15 | 1,000.00 | 0.5% | - | 0.00 | 2.5 | 0.25 | 150 | 40.00 | 1 | 150 | 100 | 2 M NaCl |
| X110 | S27 | AF17 | 1,000.00 | 0.5% | - | 0.00 | 0.8 | 0.08 | 150 | 40.00 | 1 | 150 | 100 | 2 M NaCl |
| X111 | S27 | AF17 | 1,000.00 | 0.5% | - | 0.00 | 0.8 | 0.08 | 150 | 40.00 | 1 | 200 | 100 | 2 M NaCl |
| X112 | S27 | AF17 | 1,000.00 | 0.5% | - | 0.00 | 0.8 | 0.08 | 150 | 40.00 | 1 | 350 | 100 | 2 M NaCl |
| X113 | S27 | AF17 | 1,000.00 | 0.5% | - | 0.00 | 0.8 | 0.08 | 150 | 40.00 | 1 | 450 | 100 | 2 M NaCl |
| X114 | S27 | AF17 | 4,000.00 | 0.5% | - | 0.00 | 3.2 | 0.32 | 150 | 40.00 | 1 | 150 | 100 | 2 M NaCl |
| X115 | S27 | AF16 | 1,000.00 | 2.0% | - | 0.00 | 3.2 | 0.32 | 150 | 40.00 | 1 | 150 | 100 | 2 M NaCl |
| X116 | S28 | AF18 | 1,000.00 | 0.5% | - | 0.00 | 0.8 | 0.08 | 150 | 40.00 | 1 | 150 | 100 | 2 M NaCl |
| X117 | S28 | AF18 | 1,000.00 | 0.5% | - | 0.00 | 0.8 | 0.08 | 150 | 40.00 | 1 | 200 | 100 | 2 M NaCl |
| X118 | S28 | AF18 | 1,000.00 | 0.5% | - | 0.00 | 0.8 | 0.08 | 150 | 40.00 | 1 | 350 | 100 | 2 M NaCl |
| X119 | S28 | AF18 | 1,000.00 | 0.5% | - | 0.00 | 0.8 | 0.08 | 150 | 40.00 | 1 | 450 | 100 | 2 M NaCl |
| X120 | S28 | X107 | 2.40 | 27.0% | - | 0.00 | 1.3 | 0.13 | 150 | 40.00 | 1 | 150 | 100 | 2 M NaCl |

**Supplemental Table S6:** Controlled DNA enrichment experiment results. Percentages of host and bacterial DNA before and after enrichment experiments listed in Supplemental Table S5 were estimated by qPCR using host- and bacteria-specific primers (see Supplemental Protocol). Key: ID, experiment ID (see Supplemental Table S5); PHB, percentage of host DNA before; PHA, percentage of host DNA after; PBA, percentage of bacterial DNA after; TY, total DNA yield (ng); HY, estimated host DNA yield (ng); BY, estimated bacterial DNA yield (ng).

| ID | PHB | PHA | PBA | TY | HY | BY |
| --- | --- | --- | --- | --- | --- | --- |
| X001 | 2.0% | 50.98% | 56.74% | 38.88 | 19.82 | 22.06 |
| X002 | 2.0% | 42.23% | 49.32% | 73.60 | 31.08 | 36.30 |
| X003 | 0.2% | 5.10% | 101.77% | 20.32 | 1.04 | 20.68 |
| X004 | 2.0% | 45.10% | 62.70% | 39.20 | 17.68 | 24.58 |
| X005 | 2.0% | 60.54% | 46.75% | 32.64 | 19.76 | 15.26 |
| X006 | 2.0% | 44.13% | 45.74% | 28.64 | 12.64 | 13.10 |
| X007 | 0.0% | 2.72% | 110.93% | 38.24 | 1.04 | 42.42 |
| X008 | 2.0% | 51.49% | 51.28% | 28.20 | 14.52 | 14.46 |
| X009 | 50.0% | 120.19% | 6.43% | 17.24 | 20.72 | 1.11 |
| X010 | 50.0% | 132.00% | 3.18% | 19.00 | 25.08 | 0.60 |
| X011 | 50.0% | 154.86% | 1.86% | 11.52 | 17.84 | 0.21 |
| X012 | 50.0% | 140.93% | 1.76% | 14.12 | 19.90 | 0.25 |
| X013 | 0.0% | - | - | 4.28 | - | - |
| X014 | 0.0% | - | - | 5.28 | - | - |
| X015 | 0.0% | - | - | 2.84 | - | - |
| X016 | 0.0% | - | - | 44.80 | - | - |
| X017 | 2.0% | 60.02% | 3.18% | 16.36 | 9.82 | 0.52 |
| X018 | 0.0% | - | - | 18.56 | - | - |
| X019 | 0.0% | - | - | 9.68 | - | - |
| X020 | 50.0% | 86.25% | 0.48% | 464.00 | 400.20 | 2.23 |
| X021 | 2.0% | 58.69% | 29.84% | 24.40 | 14.32 | 7.28 |
| X022 | 2.0% | 41.37% | 56.03% | 23.88 | 9.88 | 13.38 |
| X023 | 51.0% | 29.76% | 15.36% | 0.50 | 0.15 | 0.08 |
| X024 | 44.1% | 2.19% | 8.89% | 0.56 | 0.01 | 0.05 |
| X025 | 2.0% | 11.68% | 10.47% | 114.00 | 13.32 | 11.94 |
| X026 | 5.0% | 70.00% | 18.42% | 54.40 | 38.08 | 10.02 |
| X027 | 10.0% | 98.29% | 9.19% | 93.60 | 92.00 | 8.60 |
| X028 | 2.0% | 46.22% | 37.09% | 35.48 | 16.40 | 13.16 |
| X029 | 2.0% | 3.90% | 3.26% | 346.80 | 13.53 | 11.31 |
| X030 | 2.0% | 47.61% | 4.07% | 24.28 | 11.56 | 0.99 |
| X031 | 2.0% | 40.93% | 11.19% | 23.60 | 9.66 | 2.64 |
| X032 | 2.0% | 47.85% | 1.96% | 25.12 | 12.02 | 0.49 |
| X033 | 2.0% | 45.76% | 41.10% | 24.04 | 11.00 | 9.88 |
| X034 | 2.0% | 39.32% | 28.48% | 18.12 | 7.12 | 5.16 |
| X035 | 2.0% | 74.85% | 13.87% | 15.04 | 11.26 | 2.09 |
| X036 | 2.0% | 60.00% | 16.84% | 12.80 | 7.68 | 2.16 |
| X037 | 0.0% | 0.00% | 73.67% | 2.56 | 0.00 | 1.89 |
| X038 | 100.0% | 115.83% | 0.01% | 12.76 | 14.78 | 0.00 |
| X039 | 2.0% | 7.90% | 6.71% | 158.80 | 12.54 | 10.66 |
| X040 | 2.0% | 10.89% | 10.28% | 98.80 | 10.76 | 10.16 |
| X041 | 2.0% | 7.67% | 9.32% | 232.80 | 17.86 | 21.70 |
| X042 | 2.0% | 35.24% | 38.99% | 41.60 | 14.66 | 16.22 |
| X043 | 100.0% | - | - | 38.40 | - | - |
| X044 | 100.0% | - | - | 42.40 | - | - |
| X045 | 100.0% | - | - | 78.80 | - | - |
| X046 | 100.0% | - | - | 141.60 | - | - |
| X047 | 100.0% | - | - | 282.40 | - | - |

Supplemental Table S6 – continued from previous page
| ID | PHB | PHA | PBA | TY | HY | BY |
| --- | --- | --- | --- | --- | --- | --- |
| X048 | 100.0% | - | - | 456.00 | - | - |
| X049 | 100.0% | - | - | 17.52 | - | - |
| X050 | 100.0% | - | - | 14.64 | - | - |
| X051 | 100.0% | - | - | 1.89 | - | - |
| X052 | 100.0% | - | - | 3.19 | - | - |
| X053 | 0.0% | - | 2.59% | 347.20 | - | 9.00 |
| X054 | 50.0% | - | 2.19% | 496.00 | - | 10.86 |
| X055 | 0.0% | - | 10.08% | 206.40 | - | 20.80 |
| X056 | 0.5% | 0.88% | 6.01% | 220.40 | 1.95 | 13.24 |
| X057 | 2.0% | 5.32% | 3.21% | 204.40 | 10.88 | 6.56 |
| X058 | 5.0% | 12.17% | 2.88% | 222.40 | 27.06 | 6.40 |
| X059 | 0.0% | - | - | - | - | - |
| X060 | 100.0% | - | - | - | - | - |
| X061 | 2.0% | 60.57% | 19.32% | 18.36 | 11.12 | 3.55 |
| X062 | 2.0% | 83.19% | 12.39% | 14.40 | 11.98 | 1.78 |
| X063 | 2.0% | 2.10% | 0.25% | 680.00 | 14.28 | 1.70 |
| X064 | 2.0% | 1.97% | 0.30% | 656.00 | 12.92 | 1.97 |
| X065 | 2.0% | 58.82% | 18.97% | 16.32 | 9.60 | 3.10 |
| X066 | 2.0% | 101.43% | 5.27% | 9.76 | 9.90 | 0.51 |
| X067 | 2.0% | 55.65% | 16.44% | 16.28 | 9.06 | 2.68 |
| X068 | 2.0% | 65.89% | 21.69% | 17.24 | 11.36 | 3.74 |
| X069 | 2.0% | 78.68% | 17.98% | 12.76 | 10.04 | 2.29 |
| X070 | 2.0% | 128.57% | 3.58% | 6.44 | 8.28 | 0.23 |
| X071 | 2.0% | 114.15% | 2.56% | 6.36 | 7.26 | 0.16 |
| X072 | 2.0% | 28.33% | 0.82% | 26.12 | 7.40 | 0.21 |
| X073 | 2.0% | 144.68% | 4.86% | 8.64 | 12.50 | 0.42 |
| X074 | 2.0% | 133.61% | 4.28% | 7.20 | 9.62 | 0.31 |
| X075 | 2.0% | 137.91% | 6.36% | 15.88 | 21.90 | 1.01 |
| X076 | 0.5% | 237.32% | 30.14% | 2.84 | 6.74 | 0.86 |
| X077 | 0.5% | 357.50% | 24.50% | 2.40 | 8.58 | 0.59 |
| X078 | 0.5% | 270.16% | 17.74% | 2.48 | 6.70 | 0.44 |
| X079 | 0.5% | 234.81% | 33.86% | 3.16 | 7.42 | 1.07 |
| X080 | 0.5% | 257.07% | 44.62% | 3.68 | 9.46 | 1.64 |
| X081 | 0.5% | 20.03% | 2.15% | 29.16 | 5.84 | 0.63 |
| X082 | 0.0% | 0.00% | 3.59% | 26.76 | 0.00 | 0.96 |
| X083 | 2.0% | 240.00% | 8.42% | 9.00 | 21.60 | 0.76 |
| X084 | 2.0% | 243.69% | 10.49% | 8.24 | 20.08 | 0.86 |
| X085 | 2.0% | 233.55% | 20.94% | 9.36 | 21.86 | 1.96 |
| X086 | 0.5% | 227.66% | 66.81% | 3.76 | 8.56 | 2.51 |
| X087 | 0.5% | 43.31% | 12.76% | 16.44 | 7.12 | 2.10 |
| X088 | 0.0% | 0.00% | 14.38% | 14.28 | 0.00 | 2.05 |
| X089 | 0.5% | 224.26% | 45.29% | 2.72 | 6.10 | 1.23 |
| X090 | 0.5% | 46.20% | 8.26% | 16.32 | 7.54 | 1.35 |
| X091 | 2.0% | 68.14% | 3.79% | 19.40 | 13.22 | 0.74 |
| X092 | 0.0% | 0.00% | 7.89% | 15.56 | 0.00 | 1.23 |
| X093 | 0.5% | 86.44% | 7.81% | 2.64 | 2.28 | 0.21 |
| X094 | 0.5% | 73.31% | 4.84% | 2.36 | 1.73 | 0.11 |
| X095 | 0.5% | 106.31% | 3.65% | 1.74 | 1.85 | 0.06 |
| X096 | 0.5% | 114.75% | 3.60% | 1.42 | 1.63 | 0.05 |
| X097 | 0.5% | 127.97% | 40.42% | 2.36 | 3.02 | 0.95 |
| X098 | 0.5% | 130.90% | 28.13% | 2.88 | 3.77 | 0.81 |
| X099 | 2.0% | 106.34% | 8.35% | 13.56 | 14.42 | 1.13 |
| X100 | 5.0% | 135.67% | 5.32% | 17.44 | 23.66 | 0.93 |
| X101 | 0.5% | 49.11% | 40.65% | 2.48 | 1.22 | 1.01 |
| X102 | 2.0% | 83.82% | 8.76% | 13.60 | 11.40 | 1.19 |

Supplemental Table S6 – continued from previous page
| ID | PHB | PHA | PBA | TY | HY | BY |
| --- | --- | --- | --- | --- | --- | --- |
| X103 | 5.0% | 96.18% | 13.18% | 20.40 | 19.62 | 2.69 |
| X104 | 0.5% | 91.02% | 50.19% | 2.16 | 1.97 | 1.08 |
| X105 | 2.0% | 74.20% | 6.46% | 12.56 | 9.32 | 0.81 |
| X106 | 0.5% | - | - | - | - | - |
| X107 | 0.5% | 139.38% | 72.94% | 3.20 | 4.46 | 2.33 |
| X108 | 0.5% | 101.79% | 46.19% | 2.68 | 2.73 | 1.24 |
| X109 | 0.5% | 124.12% | 45.74% | 2.72 | 3.38 | 1.24 |
| X110 | 0.5% | 98.30% | 32.05% | 2.24 | 2.20 | 0.72 |
| X111 | 0.5% | 138.57% | 21.16% | 1.96 | 2.72 | 0.41 |
| X112 | 0.5% | 128.51% | 40.00% | 1.48 | 1.90 | 0.59 |
| X113 | 0.5% | 137.04% | 83.33% | 1.08 | 1.48 | 0.90 |
| X114 | 0.5% | 113.78% | 21.12% | 7.84 | 8.92 | 1.66 |
| X115 | 2.0% | 141.15% | 2.73% | 10.40 | 14.68 | 0.28 |
| X116 | 0.5% | 60.37% | 53.60% | 2.72 | 1.64 | 1.46 |
| X117 | 0.5% | 92.22% | 45.12% | 2.42 | 2.23 | 1.09 |
| X118 | 0.5% | 68.13% | 30.86% | 1.78 | 1.21 | 0.55 |
| X119 | 0.5% | 77.34% | 66.41% | 2.05 | 1.59 | 1.36 |
| X120 | 27.0% | 170.44% | 3.55% | 0.55 | 0.94 | 0.02 |

**Supplemental Table S7:**
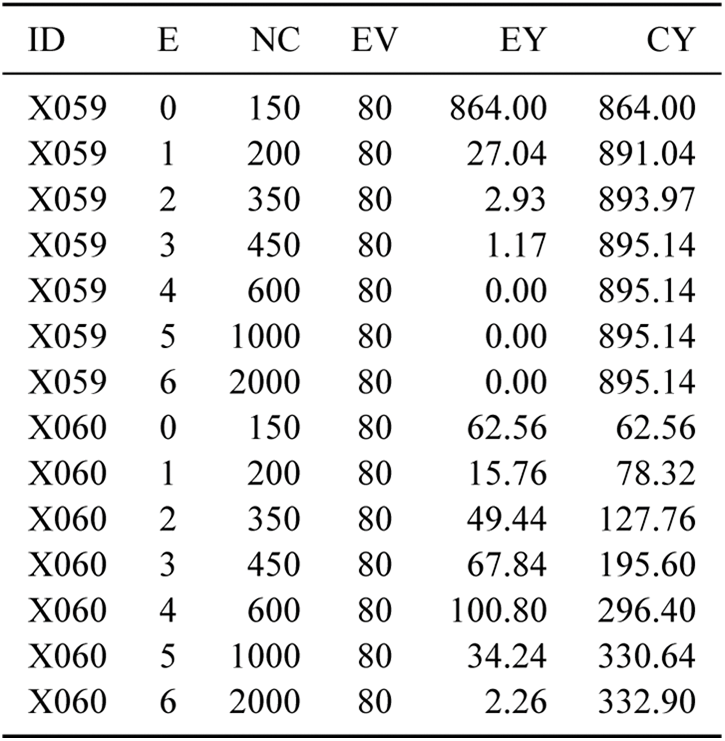
Controlled DNA enrichment elution series. After hybridizing DNA to MBD-bound beads, bound DNA was eluted in a series with progressively higher NaCl concentrations. The quantity of DNA in each elution was then quantified by Qubit. Key: ID, experiment ID (see Supplemental Table S5); EN, elution number (elution 0 represents a wash); NC, NaCl concentration of reaction (µM); EV, Elution volume (µl); EY, elution DNA yield (ng); CY, cumulative DNA yield including previous elutions in the series (ng).

| ID | E | NC | EV | EY | CY |
| --- | --- | --- | --- | --- | --- |
| X059 | 0 | 150 | 80 | 864.00 | 864.00 |
| X059 | 1 | 200 | 80 | 27.04 | 891.04 |
| X059 | 2 | 350 | 80 | 2.93 | 893.97 |
| X059 | 3 | 450 | 80 | 1.17 | 895.14 |
| X059 | 4 | 600 | 80 | 0.00 | 895.14 |
| X059 | 5 | 1000 | 80 | 0.00 | 895.14 |
| X059 | 6 | 2000 | 80 | 0.00 | 895.14 |
| X060 | 0 | 150 | 80 | 62.56 | 62.56 |
| X060 | 1 | 200 | 80 | 15.76 | 78.32 |
| X060 | 2 | 350 | 80 | 49.44 | 127.76 |
| X060 | 3 | 450 | 80 | 67.84 | 195.60 |
| X060 | 4 | 600 | 80 | 100.80 | 296.40 |
| X060 | 5 | 1000 | 80 | 34.24 | 330.64 |
| X060 | 6 | 2000 | 80 | 2.26 | 332.90 |

## Supplemental Protocol

### FecalSeq enrichment protocol

Portions of this protocol are modified from the NEBNext Microbiome DNA Enrichment Kit manual (New England Biolabs cat. #E2612S)

#### Materials and reagents

- Extracted fecal-derived DNA of known quantity
- NEBNext Microbiome DNA Enrichment Kit (New England Biolabs; cat. #E2612S or #E2612L)
- Rotating mixer
- Magnetic rack for 1.5/2.0 ml microcentrifuge tubes
- 5 M NaCl

We recommend using siliconized or LoBind tubes throughout all enrichment procedures to minimize DNA loss (Gaillard and Strauss 1998).

#### Before beginning

1. Extract and prepare DNA samples While any fecal DNA (fDNA) extraction method should in principle be compatible with the MBD enrichment, methods that maximize the recovery of host DNA are preferable. Bead-beating methods that increase total DNA yield from feces, for example, should be avoided because the mechanical disruption increases the yield of cell-wall-bound DNA (i.e., from bacteria or plants) while fragmenting host DNA. We suggest aiming for a total yield of 1 µg of DNA for all samples in a maximum volume of 30 µl each, although we have had success with as little as 500 ng (the yield of host DNA is likely more important than the yield of total fDNA). If the volume is greater than 30 µl, the DNA can be concentrated via a bead cleanup (Auxiliary protocol A). Prior to enrichment, DNA should be quantified for the total yield (e.g., by fluorometer or spectrophotometer). Ideally, the host DNA should also be quantified by qPCR (Auxiliary protocol B).
2. Calculate the required volume of MBD2-Fc-bound magnetic beads (hereafter referred to as “MBD beads”) for each enrichment reaction, as well as the total volume for a set of reactions as follows. As an approximate rule, prepare 1 µl of MBD beads for every 6.25 ng of target host DNA in each enrichment reaction. If samples contain less than 6.25 ng of host DNA or if the amount of host DNA is not quantified, prepare 1 µl of MBD beads. We recommend preparing batches of MBD beads (see step 5) with a minimum volume of 40 µl, as lower volumes preclude adequate mixing. If a smaller volume is needed, leftover unused MBD beads can be stored at 4 °C for up to a week.
3. Resuspend protein A magnetic beads by gently pipetting the mixture up and down until the suspension is homogenous, or by slowly rotating the mixture at 4 °C for 15 minutes. **Do not vortex**.
4. Prepare 1X bind/wash buffer by diluting 1 part 5X bind/wash buffer with 4 parts DNase-free water. As a general rule, the volume of 1X bind/wash buffer needed can be calculated as: 2.5 ml + 1.2 ml × [number of enrichment reactions] The amount of 1X bind/wash buffer depends on the total volume of MBD beads and the total number of enrichment reactions. MBD beads can be prepared with a maximum volume of 160 µl in a single reaction. As very small volumes (1 – 8 µl) of beads are needed for our enrichment method, a single bead preparation reaction is nearly always sufficient. If more beads are needed, increase the number of bead preparation reactions and adjust the volume of 1X bind/wash buffer accordingly. Alternatively, for volumes up to 320 µl, prepare an additional 1 ml of 1X bind/wash buffer per bead preparation reaction and add an extra wash step (see step 14). 2.5 ml of 1X bind/wash buffer are required for a single bead preparation reaction up to 160 µl. Prepare an additional 1.2 ml of 1X bind/wash buffer per enrichment reaction. This number takes into account the volume needed to prepare 2 M NaCl elution buffer in the following step. Keep 1X bind/wash buffer on ice throughout the MBD bead preparation. For the wash steps following the capture reaction, 1X bind/wash buffer can be at room temperature.
5. Prepare 2 M NaCl elution buffer by diluting 5 M NaCl with 1X bind/wash buffer. 100 µl of 2 M NaCl elution buffer are needed per enrichment reaction. 1X bind/wash buffer has a NaCl concentration of 150 mM. 1 ml of 2 M NaCl elution buffer can be prepared by adding 370 µl of 5 M NaCl with 630 µl of 1X bind/wash buffer.

#### Preparing MBD beads

1. If preparing 40 µl of MBD beads, add 4 µl of MBD2-Fc protein to 40 µl of protein A magnetic beads in a 1.5 ml microcentrifuge tube. For preparing other volumes (*n* µl) of MBD beads, add *n*/10 µl MBD2-Fc protein to *n* µl of protein A magnetic beads. As a rule, we do not prepare less than 40 µl of MBD beads due to diminished efficiency of both rotational mixing and magnetic separation at low volumes.
2. Mix the bead-protein mixture by rotating the tube in a rotating mixer for 10 minutes at room temperature.
3. Briefly spin the tube and place on the magnetic rack for 2 – 5 minutes until the beads have collected to the wall of the tube and the solution is clear.
4. Carefully remove and discard the supernatant with a pipette without disturbing the beads.
5. Add 1 ml of 1X bind/wash buffer (kept on ice) to the tube to wash the beads. Pipette up and down a few times to mix.
6. Mix the beads by rotating the tube in a rotating mixer for 3 minutes at room temperature.
7. Briefly spin the tube and place on the magnetic rack for 2 – 5 minutes until the beads have collected to the wall of the tube and the solution is clear.
8. Carefully remove and discard the supernatant with a pipette without disturbing the beads.
9. Repeat steps 10 – 13. If preparing between 160 µl and 320 µl of beads, repeat steps 10 – 13 twice for a total of three washes to ensure the removal of unbound MBD2-Fc protein.
10. Remove the tube from the rack and add *n* µl (determined in step 6) of 1X bind/wash buffer to resuspend the beads. Mix by pipetting the mixture up and down until the suspension is homogenous.

#### Capture methylated host DNA

Since reaction volumes are well under 100 µl, multiple enrichment reactions can be processed together in a microplate, with pipetting steps conducted using a multichannel pipettor. Compatible rotating mixers and magnetic separators would also be required. Here, we proceed to describe the capture procedure using a 1.5 ml tube.

The total volume of the capture reaction is an important consideration. We have observed decreased DNA binding efficiency when the concentration of MBD beads or DNA in the capture reaction is low. We therefore recommend maintaining a total reaction volume of approximately 40 µl, as we have experienced consistent success with this volume even when adding as little as 1 µl of MBD beads. Decreasing the reaction volume may result in decreased efficacy of rotational mixing. It is a good idea to keep the volume of all reactions consistent as this facilitates processing of many samples and, if DNA amounts and bead volumes are kept consistent, serves as a control for the effects of bead or DNA concentration on enrichment efficiency. Our subsequent procedures assume a reaction volume of 40 µl (not including MBD beads). If using other reaction volumes, pay particular attention to notes following each step in this section.

1. Aliquot 8 µl of 5X bind/wash buffer to a 1.5 ml microcentrifuge tube For reaction volumes other than 40 µl, tune the volume of 5X bind/wash buffer to maintain 1X concentration and adjust accordingly the volume of DNase-free water added in step 17. The volume of MBD beads should be excluded from this calculation as prepared MBD beads are already at 1X concentration. We recommend equilibrating 5X bind/wash buffer to room temperature prior to aliquoting for more accurate pipetting.
2. Add up to 30 µl of DNA (prepared in step 1) to the tube. Bring the total volume to 40 µl with DNase-free water. For reaction volumes other than 40 µl, adjust the volume of DNase-free water added to reach the target volume. Be sure to maintain 1X bind/wash concentration.
3. Add MBD beads to the tube using the volume determined in step 2. Pipette the mixture up and down or swirl a few times to mix. As an approximate rule and as stated above, add 1 µl of MBD beads for every 6.25 ng of target host DNA in each enrichment reaction. If samples contain less than 6.25 ng of host DNA or if the amount of host DNA is not quantified, add 1 µl of MBD beads.
4. Incubate the reaction for 15 minutes at room temperature with rotation.
5. Following incubation at room temperature, briefly spin the tube and place on the magnetic rack for 5 minutes until the beads have collected to the wall and the solution is clear.
6. Carefully remove the supernatant with a pipette without disturbing the beads. The supernatant is enriched for microbial DNA and may be saved and purified by bead cleanup (Auxiliary protocol A). Otherwise, discard the supernatant.
7. Add 1 ml of 1 bind/wash buffer (kept at room temperature) to wash the beads. If processing in a microplate, decrease the volume of wash buffer to 100 µl.
8. Carefully remove and discard the wash buffer with a pipette without disturbing the beads.
9. *Optional*. Add 100 µl of 1X bind/wash buffer (kept at room temperature) to the beads. Pipette the mixture up and down a few times to mix. We have found that an additional wash with 100 µl of 1X bind/wash buffer followed by rotation (steps 24 – 27) substantially improved enrichment. To skip this wash, proceed to step 28.
10. Mix the beads by rotating the tube in a rotating mixer for 3 minutes at room temperature.
11. Briefly spin the tube and place on the magnetic rack for 2 – 5 minutes until the beads have collected to the wall of the tube and the solution is clear.
12. Carefully remove and discard the supernatant with a pipette without disturbing the beads.

#### Eluting captured host DNA

The NEBNext Microbiome Enrichment Kit includes an elution protocol for captured DNA that includes digestion of DNA-bound MBD beads with proteinase K and elution with TE buffer. We have found that elution with 2 M NaCl is just as effective, is less time consuming, and conserves proteinase K. Most importantly, we have found that DNA samples eluted with 2 M NaCl and purified by bead cleanup can be further enriched in a repeat enrichment reaction. DNA samples eluted with proteinase K and TE buffer and purified by bead cleanup in contrast produced miniscule yields following a repeat enrichment reaction.

1. Add 100 µl of 2 M NaCl (prepared in step 5 and kept at room temperature) to the beads. Pipette the mixture up and down a few times to mix. If large numbers of samples are being processed, considering lowering the elution volume such that the combined volume of DNA and SPRI beads (see Auxiliary protocol A; step 1) does not exceed the capacity of microplate wells and thereby preclude the ability to parallelize bead cleanups.
2. Mix the beads by rotating the tube in a rotating mixer for 3 minutes at room temperature.
3. Briefly spin the tube and place on the magnetic rack for 2 – 5 minutes until the beads have collected to the wall of the tube and the solution is clear.
4. Carefully remove the supernatant to a fresh microcentrifuge tube and discard beads.
5. Proceed to bead cleanup to purify sample (Auxiliary protocol A).

### Auxiliary protocols

#### Auxiliary protocol A: Bead cleanup

Portions of this protocol are modified from Pacific Biosciences protocol # 001-252-177-03.

#### Materials and reagents

- Pre-washed magnetic SPRI beads, prepared following Rohland and Reich (2012)
- 70% ethanol, freshly prepared
- 1X TE buffer
- Magnetic stand
- Centrifuge

#### Procedures

1. Add 1.5X – 1.8x volume of pre-washed magnetic beads to DNA in a 1.5 ml tube. If the combined volume of beads and DNA does not exceed the capacity of the tube or well, large numbers of bead cleanups can be conducted in parallel on a microplate.
2. Mix the bead/DNA solution thoroughly by pipetting up and down several times.
3. Vortex the beads for 5 minutes.
4. Briefly spin the tube and place on the magnetic rack for 5 minutes or until the solution is clear.
5. Carefully remove and discard the supernatant without disturbing the beads.
6. Wash beads with freshly prepared 70% ethanol. Wait 1 minute, then pipette and discard the ethanol. Use a sufficient volume of 70% ethanol to completely cover the bead pellet (e.g., 100 µl for microplates and 400 µl for 1.5 ml tubes). Slowly dispense the 70% ethanol against the side of the tube opposite the beads. Do not disturb the bead pellet.
7. Repeat step 6 above.
8. Remove residual 70% ethanol and air-dry the bead pellet for 1 minute. Spin at full speed for 2 minutes in order to collect residual 70% ethanol. Then place on the magnetic rack for 30 seconds before pipetting the residual 70% ethanol and air-drying for 1 minute.
9. Resuspend the beads in 30 – 40 µl of 1X TE buffer or another suitable DNA stabilization buffer.
10. Vortex for 1 minute, then incubate for 2 minutes. Spin the sample at full speed to pellet beads. Return to the magnet and collect the supernatant in a new 1.5 ml microcentrifuge tube.
11. Following bead cleanup, quantify with a fluorometer or spectrophotometer. Validate enrichment by qPCR (Auxiliary protocol B). Enriched DNA can be sequentially enriched by repeating the enrichment protocol adding 30 µl of the enriched product to the FecalSeq enrichment protocol: step 17.

#### Auxiliary protocol B: qPCR estimation of enrichment

#### Materials and reagents

- Extracted fDNA of known quantity
- 2X SYBR Green master mix (e.g., Qiagen cat. #204143 or ThermoFisher Scientific cat. #A25780)
- Taxon-specific primers
- DNA standards For host quantification, standards can be created by performing a dilution series (i.e., 10 ng/µl, 1 ng/µl, 0.1 ng/µl, 0.01 ng/µl) of high-quality gDNA (such as blood or liver DNA) from a suitable taxon.
- qPCR instrument

#### Procedures

1. Run samples and standards at least in duplicate. We also recommend running a positive and negative control with each set of quantifications.
2. Use primers specific to the analysis a. The proportion of host DNA can be quantified by comparing qPCR results using host-specific primers to the absolute quantification estimated by some independent means (e.g., fluorometer or spectrophotometer). For our baboon DNA quantifications, we use universal mammal primers for the *MYCBP* (c-myc) gene (Morin et al. 2001):
b. Enrichment of DNA captured with MBD beads can be quantified as above using host-specific primers with enriched methylated host DNA. Alternatively, enrichment can be estimated by observing the *n*-fold decrease in quantified levels from unenriched to enriched samples using the universal 16S rRNA primer (Corless et al. 2000). 1 µl of unenriched DNA can be diluted to the concentration of the enriched sample prior to qPCR to standardize concentrations. Because MBD enrichment can in principle be biased towards densely methylated areas of the host genome, we prefer the latter method for estimating enrichment success. 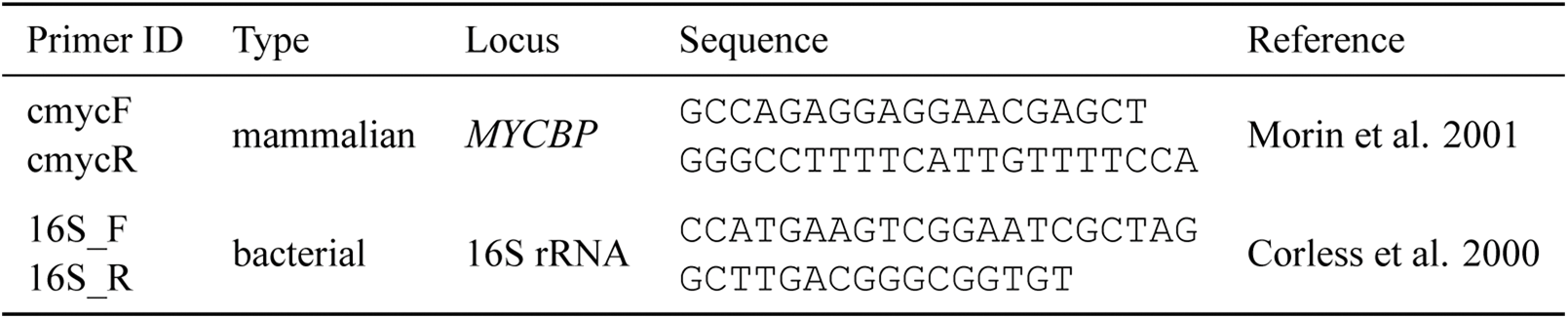
3. Set up qPCR reactions in a 20 µl total volume containing 1X of SYBR Green master mix, 0.5 mM of each primer, and 1 µl of DNA.
4. Run samples in the qPCR instrument at 95 °C for 15 minutes, followed by 50 cycles of 94 °C for 15 seconds, 59 °C (for all primers specified above; adjust for other primers) for 25 seconds, and 72 °C for 20 seconds.

## Supplemental Note

In its advertised use, the Microbiome DNA Enrichment Kit (New England Biolabs cat. #E2612S) contains enough reagents to enrich six samples, assuming 160 µll of protein A beads and 16 µl of MBD2-Fc protein are used per sample.

For FecalSeq, each reaction can be scaled down significantly. Assuming that fecal DNA samples on average contain 2.5% host DNA, we estimate that each reaction will require on average 4 µl of protein A beads and 0.4 µl of MBD2-Fc protein. This represents a scaling-down by a factor of 40. Therefore, a Microbiome DNA Enrichment Kit contains enough protein A beads and MBD2-Fc protein to support a total of 240 (6 × 40) enrichments.

240 enrichments at $168 / kit (university rate) = $0.70 per enrichment.

